# Dramatic reduction in trypanosome motility occurs without large-scale changes to paraflagellar rod ultrastructure

**DOI:** 10.1101/2024.04.19.590284

**Authors:** Heloisa B Gabriel, Ruby Kelly, Flavia Moreira-Leite, Jack D Sunter

## Abstract

Eukaryotic flagella - widely conserved structures involved in signalling, metabolism and motility – have a core microtubular axoneme that, in many organisms, is accompanied by prominent extra-axonemal structures. In kinetoplastids, including human parasites such as trypanosomes and *Leishmania*, a dense filamentous lattice called the paraflagellar rod (PFR) accompanies the axoneme for most of its length. While functional studies showed that the presence of the core PFR structure is required for normal motility, the evaluation of more subtle roles for the PFR in motility has been hampered by limited functional and localisation data, particularly on components not essential to form the ‘core’ PFR, such as signalling and metabolism proteins. Here, we addressed these issues by using the genome-wide protein localisation database TrypTag to define a PFR proteome, which was used as a base for a subtler analysis of PFR structure and function. We combined the localisation of fluorescently tagged PFR proteins relative to other cellular components with electron microscopy data on the PFR ultrastructure to localise 81 proteins to specific subdomains of the PFR. Functional analysis of a subset of PFR proteins by gene deletion and RNAi demonstrated that a novel PFR component (PFC21) is required for correct assembly of the outer PFR domain. Importantly, in some single deletion mutants, cell motility was impaired without gross disruption to the core PFR ultrastructure. Thus, our study shows that the PFR has subtle, likely regulatory roles in motility unrelated to any physical constraints that the ‘bulky’ PFR structure may impose on flagella function.

## Introduction

In eukaryotes, flagella and cilia are widely conserved organelles which extend away from the cell body and have numerous functions including motility, environmental sensing, signalling and metabolism. The core structural unit of eukaryotic flagella and cilia is the microtubule-based axoneme, which is nucleated from a basal body/centriole and is surrounded by a membrane. Many flagella also contain extra-axonemal structures, such as the fibrous sheath and outer dense fibres in mammalian sperm, vane structures in *Aduncisulus paluster* and the paraflagellar rod (PFR) in *Euglenozoa* (de Souza and Souto-Padrón, 1980; Hyams, 1982; Irons and Clermont, 1982a; Irons and Clermont, 1982b; Portman and Gull, 2010; Yubuki et al., 2016).

Extra-axonemal structures are generally substantial elements of the flagellum, with a regular, ordered appearance (Farina et al., 1986; Portman and Gull, 2010; Woolley, 1971; Yubuki et al., 2016). For example, the nine thick outer dense fibres in sperm flagella connect to each of the outer microtubule doublets, with the fibrous sheath surrounding both the axoneme and the outer dense fibres (Eddy et al., 2003; Irons and Clermont, 1982a). Together, the outer dense fibres and fibrous sheath contain many proteins involved in signalling and metabolism (Eddy et al., 2003; Petersen et al., 1999), and disruption of these structures impairs sperm motility (Miki et al., 2002; Tarnasky et al., 2010; Zhao et al., 2018). Thus, it is likely that sperm extra axonemal structures are important for flagellum beat regulation, providing both mechanical support and a structural platform for signalling and metabolism (Miki et al., 2002; Tarnasky et al., 2010; Zhao et al., 2018).

The PFR is found in most kinetoplastid parasites, including the human infective species *Trypanosoma brucei*, *Trypanosoma cruzi* and *Leishmania spp*. The flagellum of these organisms is critical throughout the life cycle, as parasites alternate between a mammalian host and an insect vector (Beneke et al., 2019; Broadhead et al., 2006; Griffiths et al., 2007). In *T. brucei*, where the PFR has been studied in more detail, the presence of a PFR is essential for motility and mammalian infectivity (Bastin et al., 1998; Griffiths et al., 2007). *T. brucei* has a long motile flagellum with a canonical ‘9+2’ axoneme (typical of motile cilia and flagella), and which extends out of a plasma membrane invagination called the flagellar pocket, near the posterior cell tip. The flagellum is laterally attached to the cell body for most of its length, via a complex of cross-linked structures that comprise the flagellum attachment zone or FAZ (Sunter and Gull, 2016). The PFR is positioned parallel to the axoneme and is tethered to axonemal outer doublets 4-7, and also to the FAZ, with the PFR first appearing as the flagellum exits the flagellar pocket, and then running along the entire length of the flagellum, until the final 750 nm of the flagellum, at which point the PFR tapers off (Höög et al., 2014). Ultrastructurally, the PFR has a ‘paracrystalline’ appearance and, in cross section, can be sub-divided into three readily identifiable domains - referred to as ‘proximal’, ‘intermediate’ and ‘distal’ - relative to the axoneme, with the proximal and distal domains containing complex layers of orderly arranged filaments (Farina et al., 1986; Portman and Gull, 2010). A recent cryo-electron tomography study highlighted the intricate, periodic nature of the structures throughout the different domains of the PFR (Zhang et al., 2021).

Earlier biochemical studies identified a number of PFR proteins including the major structural components PFR1 and PFR2, as well as calmodulin, adenylate kinases, and cAMP-specific phosphodiesterases (Bastin et al., 1998; Ginger et al., 2013; Portman and Gull, 2010; Pullen et al., 2004; Rascón et al., 2002; Shaw et al., 2019). Subsequently, a targeted approach combining proteomics with RNAi-mediated PFR2 depletion identified a further 20 PFR proteins (Portman et al., 2009). While a purified PFR fraction had no matching mass spectrometry data (Moreira-Leite et al., 1999), trypanosomatid flagella proteomes produced by mass spectrometry were likely to contain novel PFR proteins (Portman et al., 2009); however, matching localisation of novel flagellar proteins to the PFR data was still missing. The *T. brucei* genome-wide tagging project - TrypTag - has now addressed this issue (Billington et al., 2023). The TrypTag database expanded the number of proteins assigned to the PFR from ∼40 to over 150, by providing the localisation of nearly all the proteins encoded in the *T. brucei* genome, via both N– and C-terminal fluorescence tagging.

Although yeast two-hybrid analysis revealed sub-groups of protein interactions within the PFR (Lacomble et al., 2009), the specific localisation of proteins within the PFR remains largely unknown, even for the major proteins PFR1 and PFR2. Depletion and deletion studies in *T. brucei* and *L. mexicana* showed that a PFR remnant resembling the PFR proximal domain can be formed in the absence of PFR2, suggesting that PFR2 may be more concentrated in the intermediate and distal domains, while PFR1 is found throughout the PFR (Bastin et al., 1998; Bastin et al., 1999a; Maga et al., 1999; Santrich et al., 1997). This interpretation can be misleading, however, as PFR1 and PFR2 are highly similar in sequence, making it feasible that PFR1 could partially complement for the absence of PFR2, forming a PFR remnant that is structurally similar - yet biochemically different - to the proximal domain of a wild type PFR. Given the technical difficulties typically associated with immunogold labelling, particularly for a dense filamentous mesh such as the PFR, direct localisation data is available only for a few PFR proteins, including Rod1, which is found in the distal PFR domain (Woods et al., 1989), and calmodulin, which is found throughout the PFR (Ridgley et al., 2000).

Depletion of PFR1 and 2 proteins in *T. brucei* (Bastin et al., 1999a) and *L. mexicana* (Beneke et al., 2019) resulted in similar phenotypes, with marked defects in motility, including a reduction in flagellar beat frequency and directional cell movement, and the absence of productive motility. In addition, RNAi mediated depletion of calmodulin in *T. brucei* resulted in a failure to assemble the PFR including the microtubule connections, with a concomitant reduction in cell motility (Ginger et al., 2013). Aside from the modulation of flagellum beating, the PFR biochemical composition suggests that this structure acts as a platform for metabolism and signalling, an emerging theme for extra-axonemal structures (Ginger, 2005). Thus, the PFR is likely to have roles not directly related to the constraints that its bulky lattice imposes on the physical structure of the flagellum. However, as the loss of PFR proteins has either caused catastrophic changes to the PFR structure or had no discernible phenotype, it has not been possible to connect discrete structural changes in the PFR to specific functional changes in the flagellum.

Here, we have leveraged the near complete PFR proteome found in the TrypTag dataset to identify cohorts of proteins that localise to different domains of the PFR, aided by super-resolution confocal microscopy. Our extensive localisation data allowed us to map specific biochemistry and enzyme function to sub-domains of the PFR structure. Interestingly, we show that, for two PFR components (PFC22 and PFC23), single gene deletion resulted in a clear reduction in processive cell movement, without any gross structural changes to the PFR. Thus, we demonstrate here that the PFR has discrete roles in motility regulation that go beyond the imposition of a physical constraint on flagellum structure.

## Results

### *Trypanosoma brucei* PFR proteome consists of approximately 150 proteins

To investigate the function of the PFR, we used TrypTag to define a comprehensive PFR proteome. In the TrypTag database (http://tryptag.org) 161 proteins were annotated as localising to the paraflagellar rod (Billington et al., 2023). Figure 1A shows two examples of proteins with a PFR localisation, characterised by a clear signal along the flagellum, with no or little signal in the flagellar pocket region and fading towards the distal tip of the flagellum (Figure 1A). While manually inspecting the images for proteins annotated as PFR components, we identified 12 proteins that did not have the typical PFR localisation, and were annotated as localising to other flagellum associated structures. On the other hand, when inspecting the proteins annotated as axoneme in TrypTag, we identified a further 10 proteins which had the typical PFR localisation pattern. When we excluded duplicates of proteins present in gene arrays and our re-annotations, we identified a total of 151 PFR proteins (Table S1). This proteome contained many known PFR proteins (e.g. PFR1 and PFR2) that had previously been associated with the PFR through various approaches (Bastin et al., 1998; Deflorin et al., 1994; Lacomble et al., 2009; Portman et al., 2009). It should be noted that many of the proteins which localised to the PFR were also found in other regions of the cell, most commonly the cytoplasm.

**Figure 1:**
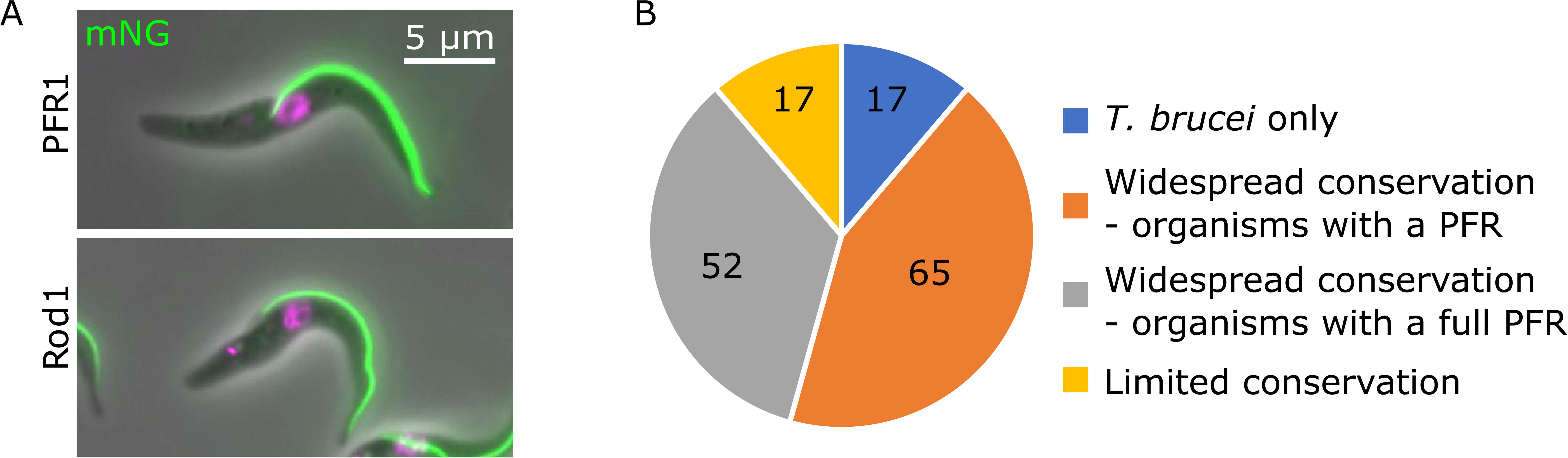
PFR proteins are well conserved across the Euglenozoa. A) Images from the TrypTag database, showing two known PFR proteins - ROD1 and PFR1 – tagged with mNeonGreen. The DNA is labelled with Hoechst (magenta). Scale bar, 5 µm. B) Pie chart showing PFR protein conservation across the Euglenozoa. The number of proteins in each group is shown.

We determined the conservation pattern of the PFR proteins across a range of eukaryotes by reciprocal best BLAST (Figure 1B, Table S1, S2). The majority (95/151) of PFR proteins were found only within the Euglenozoa, which was expected as the PFR structure is restricted to this phylum. However, those proteins with predicted signalling or metabolism function such as calpain-like and mevalonate diphosphate were also widely conserved and found in organisms outside of the Euglenozoa (Table S2). Next, we examined the conservation pattern of the PFR proteins within a large set of Euglenozoa including *Angomonas deanei* and *Strigomonas culcis*, which have a small, reduced PFR structure, and *Perkinsela sp.*, which is an endosymbiont of an amoeba and does not possess a flagellum (Fig 1B, Fig S1, Table S1). Of the 151 PFR proteins, 17 were found only in *T. brucei* and a further 17 proteins were only found in a limited number of organisms. Within this cohort of 117 conserved PFR proteins there were two groups: 65 were found in most organisms with a PFR, including those (*A. deanei* and *S. culcis*) that have a reduced PFR structure, and this set included core structural proteins (e.g. PFR1 and PFR2) and those with metabolic and signalling functions (e.g. AMP deaminase, calmodulin). A further 52 of the conserved PFR proteins were only present in organisms with a full size PFR i.e. were absent in *A. deanei* and *S. culcis* ; this group included known PFR components such as Rod1 and Par1 (Fig 1B, Table S1). Only four proteins in the PFR proteome were present in *Perkinsela sp*. - the 60S ribosomal export protein NMD3 (Tb927.7.970), AMP deaminase (Tb927.9.9740), phosphopantetheine adenylyltransferase (Tb927.11.15270) and calmodulin (Tb927.11.13020), and all of these proteins (except the common calcium binding protein calmodulin) were also found in the cytoplasm of *T. brucei*.

To understand the function of the PFR in more detail, we examined the predicted domains of the 151 PFR proteins identified with TrypTag (Table S1). The majority (∼60%) of proteins did not contain known domains; however, 11 proteins had domains associated with metabolic functions (e.g. AMP deaminase, mevalonate diphosphate) and 14 had domains with known signalling functions (e.g. calmodulin, adenylate kinase). This confirms the importance of the PFR as a platform for signalling and metabolism.

### PFR proteins can be mapped to different PFR domains by combining electron and light microscopy data

To investigate the localisation of PFR proteins within the PFR, we first examined the extensive collection of existing transmission electron microscopy (TEM) images and electron tomography datasets available in the lab, paying closer attention to the PFR configuration at the proximal (nearest the basal body) and distal regions of the flagellum. The PFR consists of three distinct domains for the majority of its length (Figure 2A, B). However, there is anecdotal evidence (backed up by TEM data in the literature and confirmed by our large collection of TEM images; Fig 2C, E) that the PFR progressively widens at its proximal end near the flagellar pocket and tapers towards the distal tip of the flagellum. To describe this detail, we examined 50 nm TEM serial sections of the proximal and distal ends of the PFR (Fig 2D, F). As we need to describe the PFR across both transverse and longitudinal axes in this manuscript, we refer to the PFR domains across the transverse axis as inner (instead of ‘proximal’), middle (instead of ‘intermediate’) and outer (instead of ‘distal’) relative to the axoneme, as the terminology ‘proximal’ and ‘distal’ is used to refer to flagellum regions closer to or further from the basal body, respectively, in the longitudinal axis.

**Figure 2:**
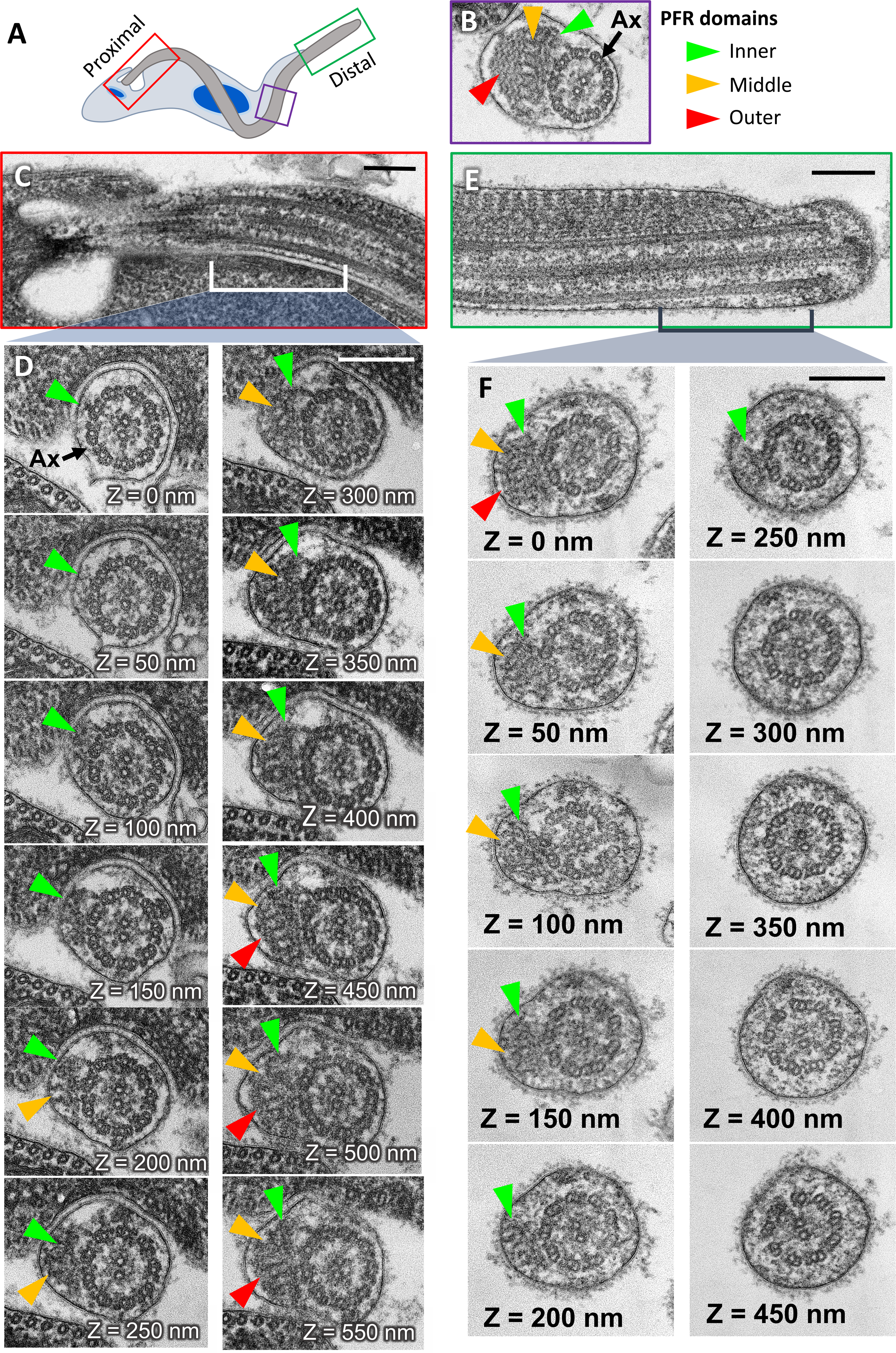
Localisation of PFR domains along the flagellum. TEM imaging of the flagellum, showing the presence / absence of PFR domains from proximal to distal. A) Cartoon of a *T. brucei* cell, with coloured boxes highlighting the proximal (red), middle (purple) and distal (green) regions of the flagellum. B) Example TEM image of a cross-section through the middle region of the flagellum, with the three domains of the PFR indicated - Inner (green arrowhead), middle (yellow arrowhead), outer (red arrowhead). Ax, axoneme. C, E) Longitudinal views of the flagellum proximal (C) and distal (E) regions, showing the appearance and progressive ‘thickening’ of the PFR as the flagellum exits the flagellar pocket (C), and the tapering of the PFR as it reaches the distal end of the flagellum (E). D, F) Serial 50-nanometer cross-sections showing the progressive appearance / disappearance of each PFR region in the proximal (D) and distal (F) ends of the flagellum. Considering the appearance of the inner domain (green arrowheads) as z = 0, the middle domain (yellow arrowheads) is first observed at ∼200 nm distal to that, followed by the outer domain (red arrowheads) which starts at ∼450 nm. The reverse is observed at the distal end (F), with the outer domain disappearing first, followed by the middle domain (lost at ∼200 nm distal from the region still containing an outer domain), and finally at ∼450 nm, the inner domain is no longer present. Scale bar, 200 nm.

The classic cross-sectional view of the PFR by TEM shows its three distinct domains, with the inner domain attached to the microtubule doublets and the middle and outer domains further away from the axoneme (Fig 2B). Our serial section analysis shows that as the PFR exits the flagellar pocket only the inner domain of the PFR was present alongside the axoneme for the first ∼150 nm. Distal to that we observed the middle domain appearing, with the inner and middle domains present together for ∼200 nm before the appearance of the clear lattice of the outer domain (Fig 2C, D). At the distal end of the PFR the reverse situation was observed, with the width of the PFR structure gradually decreasing over ∼300 nm (Fig 2E, F).

When examining the images of the tagged PFR proteins, we noticed that for many proteins the start point of the proximal end of the fluorescence signal was located either at the point of flagellum emergence from the cell body (e.g. PFR1) or slightly distal to this (e.g. Rod1) (Fig 1A). To quantify this observation, we performed a semi-automated analysis using the TrypTag dataset of the starting position of the fluorescence signal relative to the kinetoplast (i.e., the proximal end of the PFR signal along the flagellum), for 81 PFR proteins that had a strong and distinct PFR signal (Table S3). To correlate those measurements with the electron microscopy data, we subtracted the kinetoplast-to-calmodulin distance from every other measurement (Fig 3A), as calmodulin has been localised to the linkers between the axoneme and PFR (Ridgley, Webster et al. 2000) and thus is likely to be one of the most proximal PFR proteins. A group of proteins were clustered around 0 µm, extending up to ∼0.2 µm relative to the start of the calmodulin signal. Correlating this with our serial section TEM data (Fig 2C, D), we suggest this cluster contains both inner and middle PFR domain proteins. The signal of the second cluster started ∼0.5 µm relative to the calmodulin signal and this correlates with the start of the lattice structure of the outer PFR domain (∼450 nm) in our serial section analysis (Fig 2C, D). A similar analysis at the distal tip of the flagellum was inconclusive as the flagellum continues to elongate into the next cell cycle and thus the intensity and position of the fluorescence signal at the distal tip is variable (Farr and Gull 2009). Importantly, we noticed that proteins shown to interact by yeast-two hybrid, including PFR5-PFC3 and PFC4-PFC16 (Lacomble, Portman et al. 2009), were also found in the same group of ‘inner/middle’ or ‘outer’ domain proteins, based on their fluorescence signal pattern.

**Figure 3:**
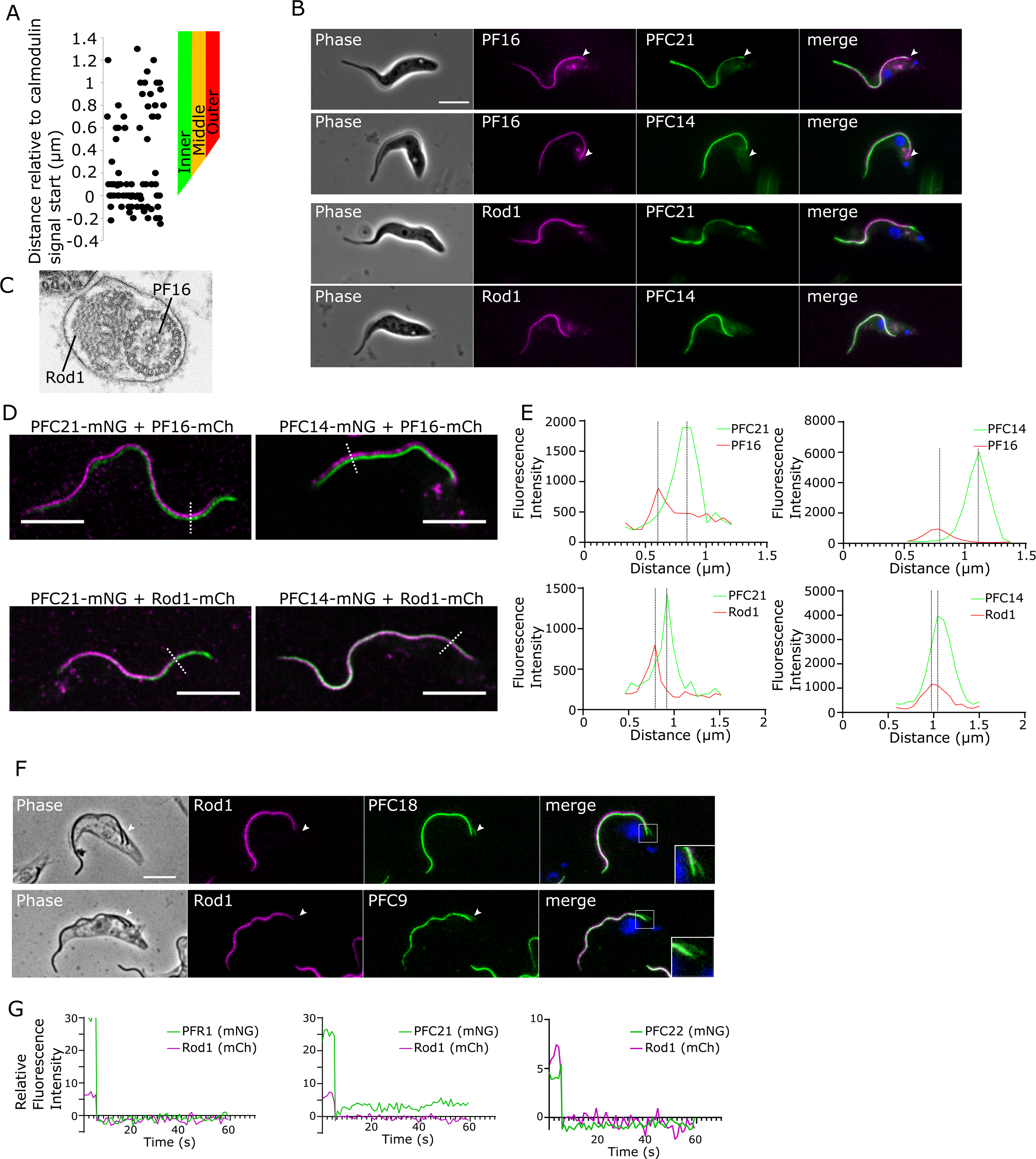
PFR protein signal ‘start point’ along the flagellum enables mapping of proteins to different PFR domains. Light microscopy analysis of PFR protein localisation, to identify patterns that can be mapped to PFR domains, as seen in TEM. A) Measurement of PFR protein signal ‘start point’ (i.e., the most proximal point of the flagellum signal) relative to the kinetoplast, for 81 mNG-tagged PFR proteins. This was a semi-automated analysis using the TrypTag dataset, where 0 represents the calmodulin signal start point (an indicator of the PFR start point; n = 15-20 cells / cell line). B) Widefield light microscopy images of cells expressing PFC21 or PFC14 endogenously tagged with mNG, with either PF16 or Rod1 endogenously tagged with mCherry. Merge shows mCherry, mNG and DNA stained with Hoechst 33342 (blue). Note that the PFC21 signal co-localises with the PF16 signal at the proximal region of the flagellum, suggesting that this protein localises to the inner/middle PFR domain (white arrowhead). In contrast, PFC14 is absent at the proximal start of the PF16 signal, suggesting localisation in the distal PFR domain (white arrowhead). Scale bar, 5 µm. C) Example TEM image of a flagellum cross-section highlighting the location of central pair protein PF16 and the outer domain protein Rod1 (cross section also shown in 2B). D) Example confocal Airyscan images of cells expressing PFC21 or PFC14 endogenously tagged with mNG, combined with either PF16 (inner axonemal marker) or Rod1 (PFR outer domain marker) endogenously tagged with mCherry. The dotted line is an example of the cross-sectioning used to measure the lateral distance between the fluorescent signals of mNG and mCherry. Scale bar, 5 µm. E) Lateral distance between signal peaks for the PFR proteins PFC21 and PFC14 (green), and either the Rod1 or PF16 markers (red), measured using confocal Airyscan microscopy. Measurements suggest that PFC21 is closer to the PF16 signal, and further from the Rod1 signal than PFC14, consistent with likely localisations in the inner/middle and outer domains, for PFC21 and PFC14, respectively. F) Cytoskeletons of early dividing cells expressing Rod1::mCherry in which PFR proteins likely to be localised in the inner/middle domain (PFC18 and PFC9) were endogenously tagged with mNG. Inner/middle domain PFR proteins are present in the new PFR (of the short new flagellum) before the outer domain protein Rod1 (arrowhead). Scale bar, 5 µm. G) Example of fluorescence recovery after photobleaching (FRAP) analysis of cells expressing Rod1 endogenously tagged with mCherry and either PFR1, PFC21, or PFC22 endogenously tagged with mNG. PFC21-mNG showed a small recovery after photobleaching.

To confirm that the proximal end of the PFR fluorescence signal was a good predictor of localisation to the ‘inner/middle’ or ‘outer’ PFR domains, we wanted to compare the position of the signal for a set of 14 PFR proteins and that of two flagellum marker proteins, the axonemal marker PF16 and Rod1, an outer PFR domain marker (Branche et al., 2006; Woodward et al., 1994). We selected mostly novel PFR proteins, with PFR1 as a control that we predicted to localise either to the inner/middle or outer PFR domains based on their fluorescence pattern, and tagged these proteins endogenously with mNeonGreen (mNG), in cell lines expressing either PF16 or Rod1 tagged with mCherry (Table 1, Fig 3C). First, we confirmed that the kinetoplast to PFR protein signal measurements in these newly generated cell lines were consistent with the data available in TrypTag (Table 1). For PFR proteins classified as inner/middle domain components, the mNG signal was closer to the kinetoplast (corresponding to 0-0.3 µm from calmodulin) than for those classified as outer PFR domain proteins (corresponding to 0.9-1.3 µm from calmodulin).

**Table 1:** Measurements of PFR protein position from kinetoplast, PF16, and Rod1. Kinetoplast to PFR protein signal start point was measured using a semi-automated algorithm in ImageJ. 15-20 cells were measured per cell line. Lateral distance measurements to PF16 and Rod1 were performed in ImageJ after confocal Airyscan processing, with inter-peak distances measured for 5-7 independent line profiles per cell line.

| Gene ID | Name | Distance to Kinetoplast ($\mu\text{m}$) | Lateral distance to PF16 ($\mu\text{m}$) | Lateral distance to Rod1 ($\mu\text{m}$) |
| --- | --- | --- | --- | --- |
| Tb927.10.9640 | PFC24 | $1.65 \pm 0.11$ | $0.15 \pm 0.02$ | $0.3 \pm 0.05$ |
| Tb927.10.8930 | PFC18 | $1.9 \pm 0.16$ | $0.16 \pm 0.04$ | $0.15 \pm 0.03$ |
| Tb927.11.14880 | PFC9 | $1.9 \pm 0.14$ | $0.17 \pm 0.05$ | $0.13 \pm 0.01$ |
| Tb927.11.13450 | PFC22 | $1.98 \pm 0.13$ | $0.18 \pm 0.02$ | $0.15 \pm 0.02$ |
| Tb927.3.4290 | PFR1 | $1.9 \pm 0.23$ | $0.21 \pm 0.02$ | $0.01 \pm 0.03$ |
| Tb927.7.6940 | PFC21 | $1.95 \pm 0.15$ | $0.25 \pm 0.04$ | $0.17 \pm 0.05$ |
| Tb927.3.5310 | PFC25 | $2.0 \pm 0.17$ | $0.24 \pm 0.04$ | $0.02 \pm 0.03$ |
| Tb927.9.9740 | AMP deaminase | $2.6 \pm 0.2$ | $0.26 \pm 0.03$ | $0.02 \pm 0.02$ |
| Tb927.3.3770 | PFC6 | $2.6 \pm 0.24$ | $0.24 \pm 0.03$ | $0.02 \pm 0.03$ |
| Tb927.10.1970 | PFC26 | $2.6 \pm 0.19$ | $0.21 \pm 0.06$ | $0.09 \pm 0.02$ |
| Tb927.7.3400 | PFC27 | $2.7 \pm 0.19$ | $0.23 \pm 0.03$ | $0.1 \pm 0.04$ |
| Tb927.10.10140 | PFC19 | $2.95 \pm 0.3$ | $0.28 \pm 0.03$ | $0.08 \pm 0.02$ |
| Tb927.10.9570 | PFC14 | $2.95 \pm 0.3$ | $0.34 \pm 0.06$ | $0.03 \pm 0.03$ |
| Tb927.4.3370 | PFC23 | $3.0 \pm 0.27$ | $0.33 \pm 0.03$ | $0.03 \pm 0.04$ |

Next, we used super-resolution (Airyscan) confocal microscopy, with a resolution limit of approximately 120 nm (Huff et al., 2017) to measure the lateral distance between the fluorescence signal peak for the mNG-labelled PFR protein and that of either Rod1 or PF16 (Fig 3D, E, Table 1). Proteins predicted to be in the inner/middle PFR domain such as PFC21 were closer to PF16 (distance between peaks = 150-240 nm) than proteins predicted to be in the outer PFR domain such as PFC14 (230 - 340 nm inter-peak distance). On the other hand, proteins predicted to be in the inner/middle PFR domain were further from Rod1 (120-300 nm inter-peak distance) than proteins predicted to be in the outer PFR domains (distance between peaks = 26-100 nm) (Table 1). However, the signals for two proteins predicted to be in the inner domain (PFR1 and PFC25) also overlapped with that of Rod1, suggesting that these proteins are present across multiple domains of the PFR.

The different domains of the PFR are likely to have different functions; we therefore mapped our bioinformatic analysis onto the PFR domains based on our protein localisation analysis. The inner/middle PFR domain contained well-conserved PFR components (e.g. PFR1, PFR2) and signalling proteins such as Protein Kinase A and calmodulin (Table S3). The outer PFR domain included proteins associated with metabolism (e.g. AMP deaminase, inositol hexakisphosphate), as well as PFR1 and other (potentially structural) PFR components such as Rod1 and PFC1. Interestingly, many of the proteins present in the outer domain of the *T. brucei* PFR were present in *S. culicis* and *A. deanei* (Table S3). One highlight of this analysis was the differential localisation of the kinases, with for example adenylate kinases located in the outer domain and the protein kinase A complex in the inner/middle domain, suggesting specialised signalling roles for the different domains.

### The PFR inner/middle domain is assembled before the outer domain

Trypanosomes are an ideal system to investigate the temporal dynamics of flagellum assembly as the old flagellum is not disassembled during the cell cycle and a new flagellum is assembled alongside the existing flagellum, at a defined position. Flagella are assembled at their distal end, with cargo transported via the intraflagellar transport system (Bastin et al., 1999b). Given the proximal position of the start of the PFR inner domain in comparison to the middle and outer PFR domains, we hypothesised that proteins in the inner/middle PFR domain would be assembled into the PFR before those in the outer domains. To assess this, we analysed cells expressing Rod1::mCherry (an outer PFR domain marker) in which PFR proteins predicted to be in the inner domain (PFC9, PFC18) were tagged with mNG, and determined the temporal order of appearance of Rod1::mCherry and PFR proteins in the new flagellum (Fig 3F). Each of these proteins predicted to be in the inner domain were present in short new flagella lacking a Rod1::mCherry signal, suggesting that these proteins are assembled into the flagellum before Rod1.

Although the PFR is a stable cytoskeletal structure (Bastin et al., 1999b), previous work has shown that there is a level of protein turnover within the PFR (Subota et al., 2014); however, in that study turnover was examined-over several hours. We reasoned that the defined PFR ultrastructure might function to enable rapid protein movement within a highly restricted framework. To test this hypothesis, we examined whether the fluorescence signal for PFR1, Rod1, PFC21 (Tb927.7.6940), and PFC22 (Tb927.11.13450) would recover after photobleaching. For PFR1, Rod1, and PFC22 there was no recovery after photobleaching (Fig 3G). However, for PFC21 we saw a small but reproducible recovery in signal, suggesting that a subpopulation of PFC21 is able to move rapidly within the PFR (Fig 3G).

### The inner/middle domain protein PFC21 is required for PFR assembly

To further dissect the role of the PFR, we investigated the function of 14 PFR proteins except PFR1 that we had mapped above. To analyse the role of these proteins on cell growth and PFR assembly, we generated gene deletions for six of these 13 proteins using CRISPR/Cas9 genome modification in a cell line expressing Rod1::mCherry. The deletion of the targeted gene was confirmed by diagnostic PCR (Fig S2). For those proteins we were unable to delete, we introduced instead a specific RNAi construct into cell lines expressing Rod1::mCherry, with the PFR protein to be depleted tagged with mNG. We confirmed successful depletion of the mNG tagged PFR proteins by fluorescence microscopy, after RNAi induction (Fig S3). Deletion or depletion of 10 of these proteins had little or no effect on cell growth or Rod1 localisation (Fig S3).

The loss of PFR2 in *T. brucei* is accompanied by the accumulation of material at the distal tip of the flagellum forming a ‘blob’, which contained PFR proteins (Bastin et al., 1999a). The deletion of either PFC21, or PFC22 caused a large reduction in cell growth (Fig 4A), and in these deletion mutants we observed a ‘blob’ at the flagellum tip, by phase contrast microscopy, in a high percentage of cells with one flagellum (Parental - ∼14%, PFC21 KO - ∼64%, PFC22 KO - ∼60%, n<20 for each) (Fig 4B). We examined the other mutants and found this blob was also present in the PFC23 deletion mutant (PFC23 KO - ∼62%, n<20) even though this null mutant did not have a large growth defect.

**Figure 4:**
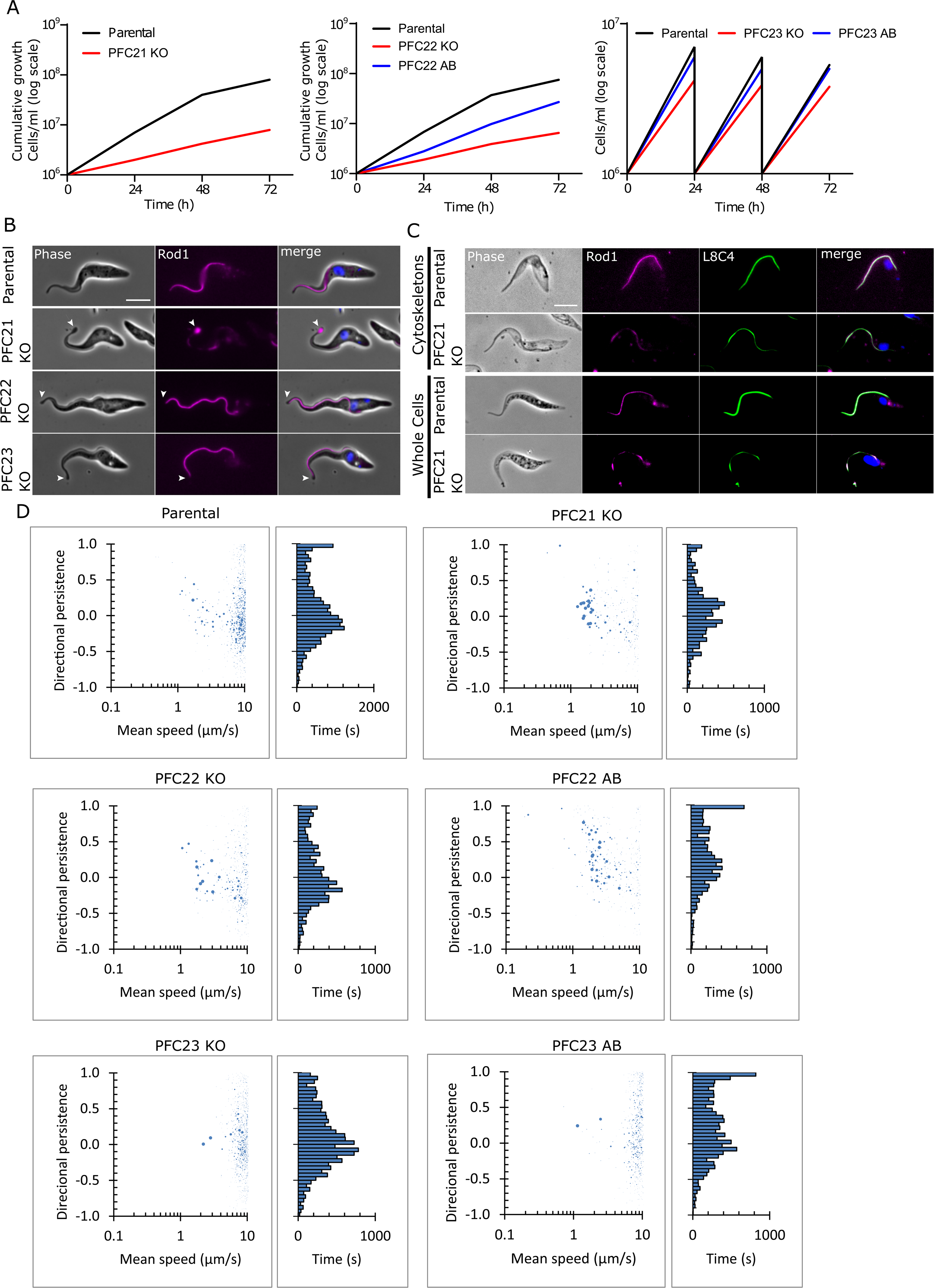
Deletion of PFC21, PFC22 and PFC23 reduced cell motility and population growth. A) Growth curves of PFC21, PFC22 and PFC23 knockout (KO) cells lines in comparison to the parental and corresponding addback (AB) cell lines. Due to the slow growth of the PFC21 and PFC22 KO cell lines, cells were seeded at 1 x 10 cells/ml and allowed to grow for 72 h. For the PFC23 cell lines, the cells were split every 24 h to 1 x 10 cells/ml and the subsequent growth followed. B) Widefield light microscopy images of live parental and knockout (PFC21, PFC22, and PFC23 KO) cells expressing Rod1 endogenously tagged with mCherry. The Rod1 signal along the flagellum is severely disrupted upon PFC21 deletion only, with Rod1 accumulating in a dilation (‘blob’) at the flagellum tip (arrowheads). PFC22 and PFC23 KO cells also have a (somewhat less conspicuous) blob, but Rod1 does not appear accumulated in the blob. The DNA is stained with Hoechst 33342 (blue). Scale bar, 5 µm. C) Widefield images of whole cells (fixed) and cytoskeletons of parental cells and PFC21 KO expressing Rod1::mCh and labelled with the anti-PFR2 monoclonal antibody L8C4. The Rod1 signal at the blob is absent from cytoskeletons, indicating that the protein is insoluble at the blob, while the Rod1 signal along the flagellum is detergent resistant. The DNA is stained with Hoechst 33342 (blue). Scale bar, 5 µm. D) Scatter plot of mean speed and directional persistence of parental cells, PFC21, PFC22 and PFC23 KO cells, and PFC22 and PFC23 AB cells, with a histogram of directional persistence. Representative graph for each cell line shown.

To determine whether the ‘blob’ contained any PFR material, we examined the localisation of Rod1::mCherry in whole cells (Fig 4B). In the parental cells Rod1::mCherry localised along the length of the flagellum with a smooth continuous signal. This pattern was seen for two of these three deletion mutants, with no clear accumulation of Rod1 in the blob. For the PFC21 null mutant the Rod1::mCherry signal along the flagellum was discontinuous with a bright signal at the tip of the flagellum associated with the ‘blob’. To investigate this phenotype in more detail, we used the monoclonal antibody L8C4 in whole cells and detergent extracted cytoskeletons to examine the distribution of PFR2 (Fig. 4C). In the parental cells, PFR2 was detected along the length of the flagellum after it had emerged from the cell body in both whole cells and cytoskeletons. While in the PFC21 null mutant the PFR2 signal was discontinuous in both the whole cells and the cytoskeletons. The ‘blob’ did not appear to survive detergent extraction, as there was no ‘blob’ at the tip of the flagellum and no associated PFR2 signal in cytoskeletons, differing to the situation observed in whole cells with a bright PFR2 signal at the tip of the flagellum associated with the ‘blob’ (Fig 4C). This suggests that PFC21 is a newly described PFR protein necessary for correct PFR assembly.

### Motility defects are not always connected with major disruptions to PFR ultrastructure

The PFR has a critical role in cell motility and to determine whether any of these mutants had motility defects, we extracted the speed and directional persistence of 1000s of cells from videomicrographs of these cells moving in media (Fig 4D) (Wheeler, 2017). For the parental cells two peaks were seen for directional persistence - one group of cells were moving in a directionally persistent manner, while the other group were moving with limited directional persistence. The movement of the PFC21, PFC22 and PFC23 null mutants was different, with the loss of the cell population moving in a consistent direction and an increase in those not moving directionally. In addition, the PFC21 and PFC22 null mutants moved slightly slower than the parental cells, while there was a small increase in the speed of the PFC23 null mutant.

To confirm the changes observed in these mutants were due to the loss of that specific gene, we generated add-back cell lines for PFC22 and PFC23 by introducing a copy of these genes tagged with mNG into their respective null mutant. We confirmed the expression of the mNG::PFC22 and mNG::PFC23 proteins by fluorescence microscopy (Fig S2). We were unable to generate add-back cell lines for PFC21. The growth of the PFC22 add-back cells was slower than that of the parental cells, but was faster than that of the deletion mutant (Fig. 4A). Moreover, the re-introduction of PFC22 into the PFC22 null mutant did not restore a parental pattern of motility of the cells (Fig 4D, Fig S4). Together, this suggests that the presence of the mNG tag may interfere with the function of the PFC22 protein. Conversely, the growth rate and the motility of the PFC23 add-back cells was similar to that of the parental cell line (Fig 4A, 4D, Fig S4). This confirms that the observed phenotype was likely due to the loss of PFC23.

To investigate whether the motility phenotype observed correlated with a defect in PFR ultrastructure, we analysed these PFR mutants by thin-section transmission electron microscopy (TEM) (Fig 5A). In the parental cells we saw the expected three domain structure of the PFR with the inner, middle and outer domains alongside the 9+2 microtubule axoneme. For PFC22 and PFC23 deletion mutants, we did not see any observable changes to the ultrastructure of the PFR, with all three domains still present and the paracrystalline structure still apparent. Conversely, in the PFC21 null mutant the outer domain was much reduced. We quantified this by measuring both the overall width of the PFR and the width of the outer domain in the TEM micrographs (Fig 5B). There was a clear reduction in the width of the PFR and specifically the outer domain in the PFC21 null mutant, suggesting that PFC21 is required for outer domain assembly.

**Figure 5:**
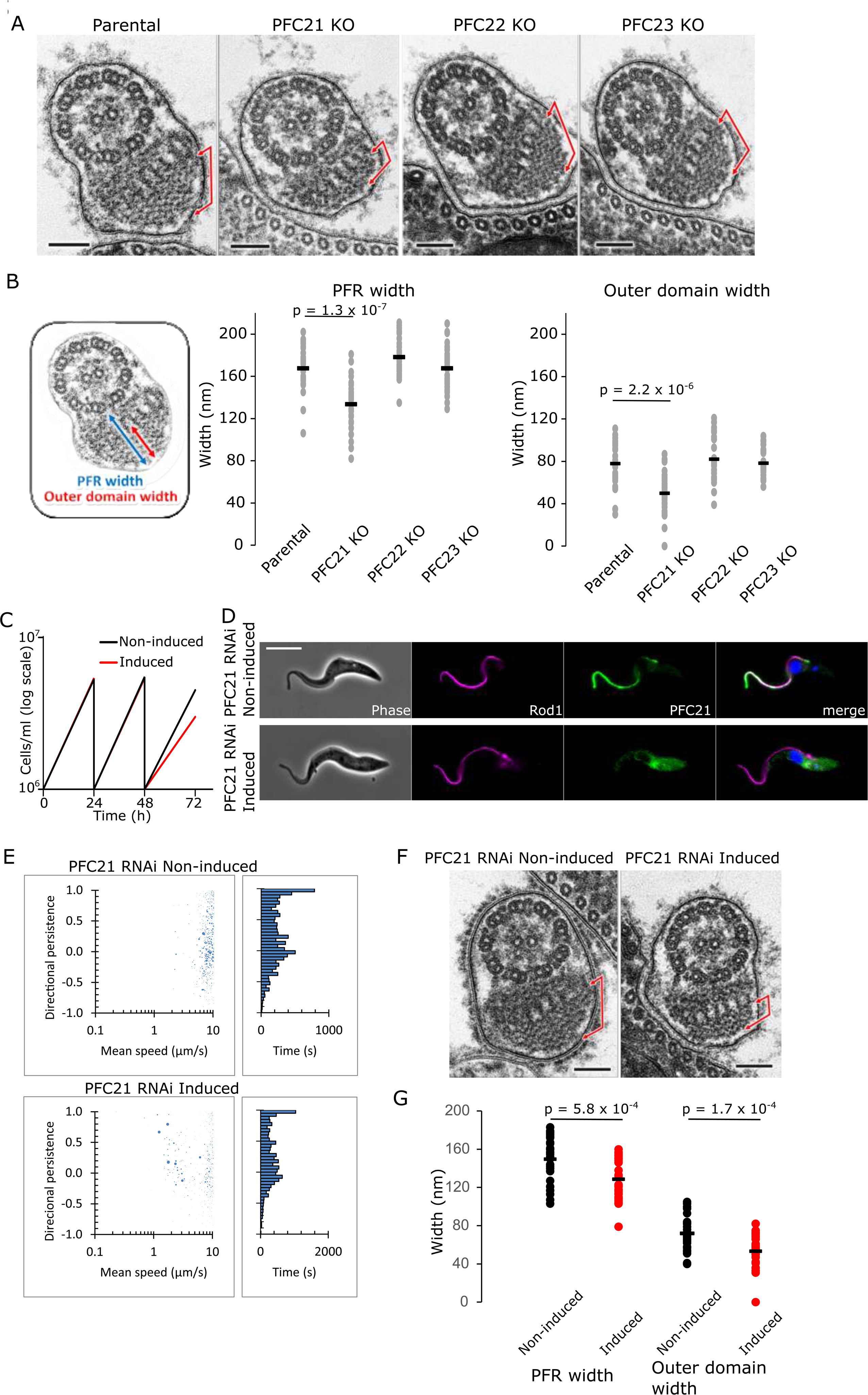
PFC21 deletion reduces the PFR outer domain, while PFC22 and PFC23 deletions have no obvious effect on the PFR ultrastructure. A, B) TEM analysis of selected PFC knockout mutants with reduced motility. A) Example TEM cross-sections of flagella from parental, PFC21, PFC22 and PFC23 KO cells. Scale bar, 100 nm. The brackets indicate the outer domain of the PFR, which is clearly reduced in PFC21 KO cells only. B) Measurement of entire PFR width and outer domain width from parental, PFC21, PFC22 and PFC23 KO cells. The diagram on the left shows the position of width measurements in an example image. Dots represent single measurements and the black line is the mean (p-value by unpaired, two-tail t-test, n=30). C – G) PFC21 RNAi knockdown analysis. C) Growth curve of PFC21 RNAi cells in the presence or absence of RNAi induction (with doxycycline) for 72 h. Cultures were split every 24 h to 1 x 10 cells/ml, to maintain exponential growth for 72 h. D) Fluorescence microscopy analysis of PFC21 RNAi induction, in cells expressing Rod1 endogenously tagged with mCherry (magenta) and PFC21 endogenously tagged with mNG (green), to monitor RNAi mediated protein knockdown. The DNA is stained with Hoechst 33342 (blue). Note that the PFC21 signal is undetected in the flagellum, after 72 h of induction. E) Scatter plot of mean speed and directional persistence of non-induced and induced (72 h) PFC21 RNAi cells, with a histogram of directional persistence. Representative graph for each cell line shown. F) Example TEM cross-sections of flagella from non-induced and induced (72 h) PFC21 RNAi cells. Scale bar, 100 nm. The brackets indicate the outer domain of the PFR, which is reduced after PFC21 RNAi. F) Measurement of entire PFR width and outer domain width (as indicated in B) from non-induced and induced (72 h) PFC21 RNAi cells. Dots represent single measurements and the black line is the mean (p-value by unpaired, two-tail t-test, n=30).

Given that we were unable to complement the deletion of PFC21, we used RNAi to confirm the role of PFC21 in PFR assembly. We analysed the effect of PFC21 depletion by introducing an inducible PFC21 RNAi construct into the cell line expressing Rod1::mCherry and PFC21::mNG. Successful depletion of PFC21 was confirmed by fluorescence microscopy (Fig 5C). Loss of PFC21 by RNAi did not have such a dramatic effect on cell growth as seen with the deletion cell line and there was little disruption of Rod1 localisation (Fig 5C, D). Moreover, we examined the effect of PFC21 depletion on cell motility and found little change in motility (Fig 5D, Fig S4). However, when we examined the PFC21 RNAi cells after 72h of induction, we observed that in many cross sections the width of the outer PFR domain was reduced as we saw in the PFC21 null mutant, providing further evidence that PFC21 is important for outer PFR domain assembly (Fig 5E, F).

## Discussion

Here, we have taken advantage of the comprehensive PFR proteome determined by TrypTag to investigate the functions of the PFR (Billington et al., 2023). We refined the initial TrypTag dataset, giving a total of 151 PFR proteins. Unsurprisingly, the majority of these proteins are restricted to the Euglenozoa; however, within the PFR cohort there were signalling and metabolic proteins with a widespread distribution. This further confirms previous work suggesting that the PFR acts as a hub for signalling/metabolism and importantly, given the genome scale of TrypTag, we now have a near complete picture of the potential pathways in the PFR (Moran et al., 2014; Portman and Gull, 2010). Yet, it should be noted that some proteins are refractory to tagging or will mis-localise when tagged with a fluorescent protein, so it is likely there are a handful of PFR proteins still to be identified. However, this study confirms that TrypTag is an invaluable resource for the analysis of organelle structure and function.

At the proximal end of the flagellum not all three domains of the PFR are present, with initially only the inner domain present. The middle and outer domains are progressively assembled as the PFR extends along the flagellum. Interestingly, when we examined the proximal start point of the PFR signal for our PFR cohort we could readily define two sets of proteins, those with a start point closer to the kinetoplast and those with a start point further away from the kinetoplast. Together this suggests that the position of the PFR protein signal start point indicates the domain in which that protein is found. We validated this observation by measuring the lateral offset between a selected set of PFR proteins and markers for the central pair and the PFR outer domain. The proteins with a start point closer to kinetoplast were found to be positioned closer to the central pair, while those with a start point further from the kinetoplast were found closer to the PFR outer domain. Our results show that many PFR proteins are not evenly distributed across the structure and are in fact present in specific domains, suggesting that each of these domains will contribute to different functional aspects of the PFR. By combining our data, including light and electron microscopy analysis, we generated a map of the localisation of PFR proteins within the different PFR domains (Fig 6).

**Figure 6:**
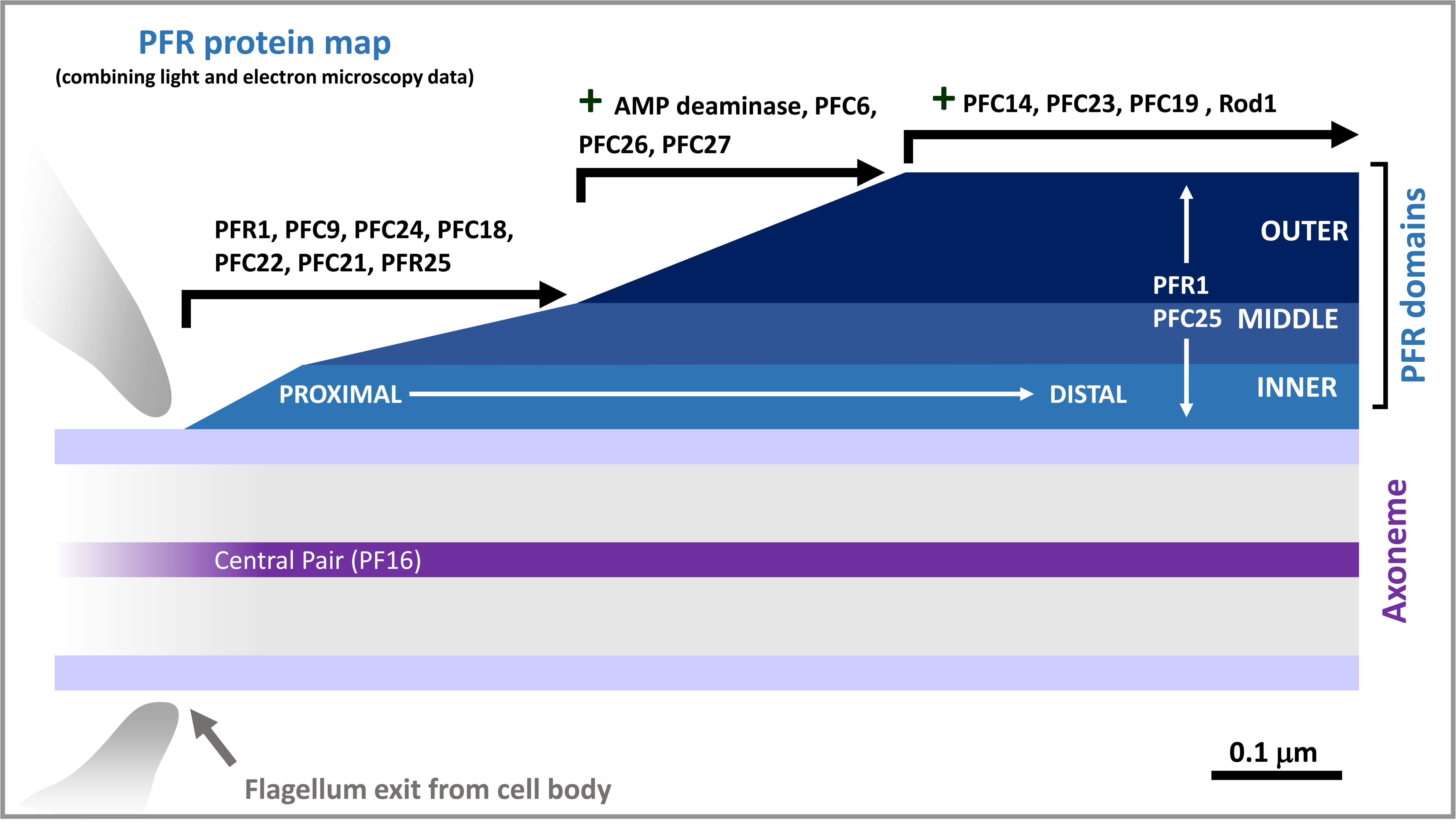
PFR protein map. Predicted localisation of PFR proteins to the different PFR domains, based on mapping of our combined light microscopy data to the known locations of inner, middle and outer PFR domains, as seen by TEM.

We extended this analysis and by examining the position of the start point of the fluorescent signal of the tagged protein relative to the kinetoplast, we have mapped 81 proteins to the inner/middle and outer PFR domains. Reassuringly, previously identified interacting PFR proteins such as PFR5-PFC3 and PFC4-PFC16 were predicted to localise to the same PFR domain as each other (Lacomble et al., 2009; Portman et al., 2009). However, this approach has an important caveat as for the majority of the proteins we are not able to define if they are solely present in specific domains and this issue is more likely to affect those proteins assigned to the inner/middle domain in this study. For example, though calmodulin was mapped to the inner/middle domain, previous work using immuno-EM showed it was present across the entire PFR (Ridgley et al., 2000). Indeed, our analysis of the lateral position of PFR1 and PFC25 suggested these proteins were present across all the different domains, as was expected for PFR1 (Portman and Gull, 2010). While bearing this caveat in mind our analysis identified important patterns of protein localisation.

Protein phosphorylation machinery was found throughout the PFR. The inner/middle PFR domain cohort contained two kinases and a cAMP-dependent protein kinase regulatory subunit, with catalytic subunits 1 and 2. Intriguingly, another cAMP-dependent protein kinase regulatory subunit (Tb927.9.10890) was found in the axoneme by TrypTag, and this differential localisation of cAMP-dependent protein kinase subunits between the PFR and axoneme potentially has a role in regulating flagellum beating. Moreover, two adenylate kinases were predicted to be in the outer PFR domains in addition to a pseudokinase and a potential protein tyrosine phosphatase. Thus, while phosphorylation signalling cascades are clearly present across the PFR, we are now able to resolve the specific domains in which the kinases/phosphatases operate.

Calmodulin was identified as an inner/middle PFR domain protein, which aligns with previous work showing calmodulin to be a component of the connections between the PFR and the axonemal microtubule doublets, in addition to being distributed across the rest of the PFR (Ginger et al., 2013; Ridgley et al., 2000). It is therefore likely that other components of these connections will be found in our inner PFR domain cohort. *A. deanei* and *S. culicis* have a reduced PFR, with ultrastructure analysis showing that a structure reminiscent of the inner PFR domain is present. However, many of the proteins predicted to be within the outer PFR domain were present in the genomes of these organisms, such as adenylate kinase, and it will be interesting to determine where in the reduced PFR these proteins are located in these parasites or if they are even located within the PFR. Moreover, this suggests that the PFR molecular organisation may be significantly different in these organisms, with the reduced PFR incorporating proteins present across all three domains of the *T. brucei* PFR.

Early work showed that the PFR was assembled at its distal end, with a level of turnover along its length (Bastin et al., 1999b). By defining inner/middle and outer PFR domain cohorts, we were able to show that inner/middle PFR domain proteins were present in the flagellum before outer PFR domain proteins. This suggests that the PFR is built with a defined assembly order extending from the axoneme out, with the connections to the microtubule doublets likely assembled first, followed by the inner domain and then the middle and outer domains. However, the marker(s) that define the position of the start point of PFR assembly remain an open question. The PFR is a key component of the trypanosome flagellum cytoskeleton, which is generally considered to be a relatively static structure, with a slow turnover of components. In terms of the PFR, a small amount of tagged PFR2 is slowly (within hours) incorporated into the old flagellum PFR that was assembled in the previous cell cycle, showing that there is limited protein turnover over a longer timescale (Bastin et al., 1999b). The majority of the proteins we examined by photobleaching did not show any recovery in signal, suggesting that these are stably integrated components of the PFR. Intriguingly, PFC21 behaved differently, with a small but rapid (within seconds) recovery in signal after photobleaching, likely due to a small pool of PFC21 being able to rapidly diffuse through the PFR. This suggests that the PFR provides a stable cytoskeletal framework within which selected components are able to rapidly move around.

To dissect the role of the different PFR domains, we screened and analysed the effect of the deletion/depletion of 15 conserved proteins drawn from both the inner/middle domain and outer domain. This highlighted the importance of three proteins - PFC21, PFC22, and PFC23 - for motility, with the loss of these proteins resulting in a drop in important motility parameters. The PFR has a well-recognised role in regulating flagellum beat and cell motility, with the depletion of PFR2 in *T. brucei* and the deletion of PFR1 and/or PFR2 in *L. mexicana* reducing the swimming capability of these mutants (Bastin et al., 1998; Maga et al., 1999; Santrich et al., 1997). The loss of PFR1 and PFR2 was accompanied by either a complete or near-complete loss of the PFR depending on the mutant; however, this severe disruption makes it difficult to pinpoint the specific role or roles of the PFR in flagellum beat control and cell motility. The PFR has therefore been suggested to have both a physical function in flagellum beating (acting as a mechanical spring), and also a regulatory function in this process (Hughes et al., 2012; Moran et al., 2014).

Our results suggest that the outer PFR domain has a key role in cell motility, as the loss of PFC21 resulted in a reduction in the PFR outer domain, as shown by the disruption to Rod1 localisation and our TEM analysis, causing a reduction in cell motility. In addition, PFC21 deletion also led to the disruption of PFR2 localisation, suggesting that PFR2 is also found within the outer domain. This agrees with earlier work in which the depletion of PFR2 in *T. brucei* resulted in a much-reduced PFR structure that resembled the inner PFR domain and a similar result was also seen when PFR2 was deleted in *L. mexicana*, suggesting that assembly of the inner domain did not require PFR2, as it was likely not located within this domain (Bastin et al., 1998; Maga et al., 1999; Santrich et al., 1997). We assessed the function of PFC21 by using both a deletion and RNAi mutant. The deletion of PFC21 caused a more severe phenotype, with a decrease in cell growth and directional persistence that was not observed for the RNAi mutant. However, in both the deletion and RNAi mutants the outer PFR domain was significantly reduced, confirming that PFC21 is necessary for complete assembly of the outer domain. The differences between the deletion and the RNAi phenotypes for PFC21 are likely related to the lower penetrance of the RNAi knockdown, where a residual amount of PFC21 is likely to be present even after 72h of induction. PFC21 and PFC22 localised to the inner PFR domains, while PFC23 localised to the middle/outer PFR domains, yet the deletion of any of these proteins resulted in defects in cell motility. This shows that all the PFR domains contribute to the role of the PFR in cell motility, and highlights the biochemical complexity required to maintain a coordinated flagellar beat.

Interestingly, the PFC22 and PFC23 null mutants had reduced directional persistence, yet the PFR ultrastructure appeared unchanged, and both Rod1 and PFR2 exhibited their usual localisation along the length of the flagellum. However, we cannot exclude the possibility of ultrastructural changes that are too subtle to be seen by our approach, especially as a ‘blob’ was seen at the flagellum tip in these mutants; in other mutants, the blob arises from a disruption in flagellum assembly. Thus, changes in cell motility can be caused by small subtle differences in PFR structure and not just the wholesale disruption seen with the loss of PFC21, calmodulin, PFR1, or PFR2 (Bastin et al., 1998; Ginger et al., 2013; Maga et al., 1999; Santrich et al., 1997). This suggests that the PFR has a key role in motility as a platform for signalling and metabolism, and not simply by providing mechanical resistance due to its large physical structure (Hughes et al., 2012). Extra-axonemal structures are common in many flagellated/ciliated organisms but are not ubiquitous, suggesting that the coordinated axoneme movement can readily be maintained with or without such additional structures. Moreover, these structures take a variety of forms and therefore it may not be the actual ‘physicality’ of the extra-axonemal structure, which is important, but rather the positioning of components/functions relative to the axoneme.

Our near complete PFR proteome of 151 proteins suggests that this structure has a role in multiple functions including metabolism, homeostasis and sensing. We have defined which PFR domain 81 of these proteins are likely to be found in, providing spatial resolution to specific signalling and metabolism pathways, with the adenylate kinases important for cAMP signalling restricted to the PFR outer domain. Moreover, we have demonstrated for the first time that motility defects can be the result of subtle changes to the PFR and not just large disruptions of the structure. This supports the idea that the intricate PFR ultrastructure acts to position and organise protein function, because the loss of specific proteins can cause functional disruption to motility on a similar scale to that of wholesale loss of the PFR structure.

## Materials and Methods

### Reciprocal Best Blast

For the reciprocal best Blast analysis, we adapted the protocol from Schmelling et. al. 2017 (Schmelling et al., 2017). The coding sequences of all entries in the GenBank protein database (Benson et al., 2008), which were labelled as “Complete Genome” or “Chromosome”, were downloaded from the NCBI FTP server (version 2021). These sequences, including the coding sequences of *Trypanosoma brucei* TREU927, were used to construct a custom protein database for the homology search. In addition, protein sequences of the 156 related proteins (Table S1), from *Trypanosoma brucei* TREU927 were used as queries for a search of homologs within the custom Euglenozoa or General protein database, applying the standalone version of BLASTP 2.10.0+ with standard parameter (wordsize: 3, substitution matrix: BLOSUM62). The 10,000 best hits with an e-value of 10^−5^ or lower were filtered for further analyses. The returned hits were used as queries for a second reverse BLASTP run, searching for homologs in *Trypanosoma brucei* TREU927 genomes using the same parameters as above with an altered e-value of 10. Only hits with the original query protein as best reversal hit were accepted for further analyses, thus minimizing false positive results. Python (version 3) was used for numerical operations, data visualization and heatmap plotting.

### Culture and cell line generation

TREU927 1339 procyclic *T. brucei* were used for all experiments (Alves et al., 2020). Cells were cultured in SDM79 media supplemented with Haemin and 10% foetal calf serum (Brun and Schönenberger, 1979) at 28°C and maintained at a density between 1 × 10 to 1 × 10 cells/ml. Analysis of growth was performed at a starting density of 2 × 10 cells/ml and monitored daily using a Z2 Coulter Counter.

Cell lines expressing PF16 (Tb927.1.2670) or Rod1 (Tb927.11.15100) endogenously tagged with mCherry as an axoneme and PFR outer domain markers respectively were generated using the PCR tagging methodology (Dean et al., 2015). Endogenous tagging of 15 selected PFR proteins with mNG was performed, as previously described, using a tagging construct encoding a 51 Ty::mNG epitope tag [28] and hygromycin resistance marker amplified from the pPOTv6 plasmid using long primers incorporating 80 nt of homology (Dean et al., 2015). For the generation of null mutants using CRISPR/Cas9 mediated genome editing, the 1339 cell line was transfected with guide and repair constructs generated by PCR using primers designed on the LeishGedit website using the G00 primer and the pPOTv6Blast and pPOTv6Hygro plasmids as templates (Beneke et al., 2017). To produce the add-back cell line, the open reading frames were amplified by PCR and cloned into the XbaI and BamHI restriction sites of the constitutive expression plasmid (pJ1313) described in (Alves et al., 2020).

The RNAi plasmids for RNAi knockdown were generated using the pQuadra system (Inoue et al., 2005). Primers were designed for the amplification of genes of interest using RNAit (Redmond et al., 2003). The PCR products and the plasmids pQuadra 1 and pQuadra 3 were digested with BstXI. Then, the PCR product and the fragment released from pQuadra 1 were ligated into the pQuadra 3 backbone. Following sequencing confirmation, the plasmid was linearized with NotI before transfection. Doxycycline (1 µg/ml) was added to cultures to induce knockdown of expression. All **c**onstructs were transfected using a Nucleofector 2b as described previously (Dean et al., 2015).

### Fixed and live-cell microscopy

For whole-cell immunofluorescence, cells were washed with PBS and allowed to settle on poly-L-lysine–coated slides before fixing with −20°C methanol for 20 min. For cytoskeleton preparations, 0.5% Nonidet P-40 in PEME (100 mM PIPES-NaOH, pH 6.9, 2 mM EGTA, 1 mM MgSO_4_, and 100 nM EDTA) was added to the cells for 30 s, followed by fixation in −20°C methanol for 20 min. For immunofluorescence, slides were incubated with mouse monoclonal antibody L8C4 (1:200) (Kohl et al., 1999) diluted in PBS with 1% BSA for 1 h at room temperature. Then, slides were washed with PBS and AlexaFluor488 conjugated goat anti-mouse IgG (H+L) cross-adsorbed secondary antibody, diluted (1:200) in PBS with 1% BSA were added to the slides, and incubated for 45 min at room temperature. Slides were then washed with PBS and stained with Hoechst 33342.

For live-cell microscopy, cells were washed in PBS supplemented with 10 mM glucose and 46 mM sucrose (vPBS). In the second wash, DNA was stained using 10 µg/ml Hoechst 33342. After the third wash, cells were resuspended in PBS, settled onto glass slides, mounted with a coverslip, and imaged immediately.

Cells and cytoskeletons were imaged using a Zeiss ImagerZ2 widefield microscope with 63x or 100x objective and Hamamatsu Flash 4 camera. Alternatively, laser confocal Airyscan Imaging was performed on a Zeiss LSM880 equipped with an Airyscan detector, using a Plan-Apochromat 63x objective. For multi-track imaging, GFP was imaged with 488 nm excitation and RFP was imaged with 561 nm excitation. Emission wavelengths were filtered through an LP460 filter. FIJI was used for contrast and brightness corrections on images, as well as for length/width measurements from light microscopy images (Schindelin et al., 2012). The distance from the kinetoplast to PFR protein signal start point was measured using a semi-automated algorithm in FIJI. A total of 15-20 cells were measured per cell line. Lateral distance measurements to PF16 and Rod1 were also performed in FIJI after Airyscan processing, with inter-peak distances measured for 5-7 independent line profiles per cell line.

Fluorescence Recovery After Photobleaching (FRAP) experiments were performed using a Zeiss LSM800 confocal microscope and Plan-Apochromat 63x objective. Pixel dwell time was 0.48 µs, pixel size was 0.05 µm and the image area was 28.35 µm x 28.35 µm. Laser powers of 1% were used for 488 nm and 568 nm lasers, with detection ranges of 400-559 nm and 589-700 nm for PFC proteins tagged with mNeonGreen and Rod1-mCherry, respectively. Cells were imaged for 5 seconds (6 frames) before bleaching. Regions along the flagellum were chosen such that anterograde and retrograde recovery could be studied, and photobleached using a 488 nm laser at 100% intensity until signal intensity dropped to 10% compared to pre-bleaching. Individual cells were then imaged for a further 55 seconds (55 frames) to record any recovery.

For cell swimming analysis, a 64 s video of 388 frames each were captured under darkfield illumination using a 10x objective. Particle tracks were traced and quantified (mean speed and cell directionality – i.e. ratio of velocity to speed) automatically as previously described (Wheeler, 2017).

### Transmission electron microscopy (TEM)

For TEM, cultures in the mid-log phase of growth were fixed in medium by the addition of glutaraldehyde for a final concentration of 2.5%. After brief (2-3 min) fixation in medium, cultures were spun at 800 g for 5 min and cell pellets were resuspended gently in a fixative solution consisting of 2.5% glutaraldehyde, 2% formaldehyde and 1% tannic acid in 0.1 M phosphate buffer (pH 7.2). Cells were incubated in this primary fixative for 1h at RT. Then, samples were washed 3x in 0.1 M phosphate buffer and post-fixed in 1% osmium tetroxide, 1.5% potassium ferricyanide in 0.1 M phosphate buffer, for 1 h at RT (and in the dark). After post-fixation, samples were washed 3x in ddH₂O and embedded (as a pellet) in low melting point agarose. Agarose-embedded cell pellets were cut into small fragments of ∼1 mm³ and stained overnight (at 4⁰C) with 1% aqueous uranyl acetate. Samples were then washed 3x in ddH₂O, dehydrated in ethanol (30%, 50%, 70%, 90% and 3x 100% ethanol, 10 min per step), and the infiltrated and embedded in epoxy resin (TAAB 812 Hard premix, TAAB, cat. no. T030). Ultrathin sections of 70 nm were obtained using an RMC PowerTome ultramicrotome, post-stained with Reynolds’ lead citrate for 5 min, and then imaged at 120kV in a JEOL 1400 Flash transmission electron microscope equipped with a OneView 16-megapixel camera (Gatan – Ametek). To make cross-section views of the flagellar cytoskeleton more comparable across samples, flagellum cross-sections were tilted up to ±25⁰ until a clear view of the protofilaments of axonemal microtubules was obtained. PFR and PFR domain width measurements were performed in ImageJ, with 30 cross-sections measured for each cell line and/or condition.

## Acknowledgements

HBG was supported by a Newton International Fellowship from the Royal Society. Work in the Sunter lab is funded by the Wellcome Trust and Leverhulme Trust. We thank Richard Wheeler for assistance with ImageJ scripts and the Oxford Brookes Bioimaging Centre for their excellent technical support and expertise, especially Ed Rea.

## Supplemental information

**Figure S1.** Heatmap of the reciprocal best BLAST analysis results. The colour indicates the percent identity between the *T. brucei* PFR protein and the ortholog in that species. White means no ortholog present.

**Figure S2.** A) PCR analysis of knockout mutants. Loss of deleted gene analysed using gene specific primers. Gene specific primers were tested against parental genomic DNA. Negative control was water only. B) Example images of PFC22 and PFC23 add back cells expressing Rod1 endogenously tagged with mCherry and PFC22 and PFC23 tagged with mNG respectively. Far left is a phase image, followed by the Rod1 marker and then PFC22 or PFC23. Merge shows mCherry, mNG and DNA stained with Hoechst 33342. Scale is 5 µm.

**Figure S3.** Example images of PFC19 and AMP deaminase deletion mutants expressing Rod1 tagged with mCherry. Left is a phase image, followed by the Rod1 marker. Merge shows phase, mCherry and DNA stained with Hoechst 33342 (blue). Scale is 5 µm. Growth of parental and KO cells was followed for 72 h. Example images of cells before and after RNAi induction for 72 hours with doxycycline (1 µg/ml). Cells express Rod1 endogenously tagged with mCherry and PFR protein of interest tagged with mNG, to monitor RNAi mediated protein knockdown. Far left is a phase image, followed by the Rod1 marker and then the PFR protein. Merge shows phase, mCherry, mNG and DNA stained with Hoechst 33342 (blue). Growth of non-induced and induced cells was followed for 72 hours.

**Figure S4.** Motility analysis of PFC mutant cell lines. Box and whisker plots of mean speed and mean directional persistence showing the minimum value, first quartile, median, third quartile and maximum value. The black X is the mean value. Each individual motility assay measures the speed and persistence of >700 cells. The mean from each of these assays for speed and directional persistence is then calculated and plotted here. For each cell line the motility assay was performed 3-6 times. P-value was calculated using a 2-tailed t-test.

**Table S1.** Results from Reciprocal Best BLAST analysis

**Table S2.** Summary from Reciprocal Best BLAST analysis

**Table S3.** Semi-automated analysis using the TrypTag dataset of the start position of the fluorescence signal relative to the kinetoplast of 81 PFR proteins.

## References

Alves, A. A., Gabriel, H. B., Bezerra, M. J. R., de Souza, W., Vaughan, S., Cunha-E-Silva, N. L. and Sunter, J. D. (2020). Control of assembly of extra-axonemal structures: the paraflagellar rod of trypanosomes. J Cell Sci 133, jcs242271.

Bastin, P., Sherwin, T. and Gull, K. (1998). Paraflagellar rod is vital for trypanosome motility. Nature 391, 548.

Bastin, P., Pullen, T. J., Sherwin, T. and Gull, K. (1999a). Protein transport and flagellum assembly dynamics revealed by analysis of the paralysed trypanosome mutant snl-1. J. Cell. Sci. 112 (Pt 21), 3769–3777.

Bastin, P., MacRae, T. H., Francis, S. B., Matthews, K. R. and Gull, K. (1999b). Flagellar morphogenesis: protein targeting and assembly in the paraflagellar rod of trypanosomes. Mol. Cell. Biol. 19, 8191–8200.

Beneke, T., Madden, R., Makin, L., Valli, J., Sunter, J. and Gluenz, E. (2017). A CRISPR Cas9 high-throughput genome editing toolkit for kinetoplastids. R Soc Open Sci 4, 170095.

Beneke, T., Demay, F., Hookway, E., Ashman, N., Jeffery, H., Smith, J., Valli, J., Becvar, T., Myskova, J., Lestinova, T., et al. (2019). Genetic dissection of a Leishmania flagellar proteome demonstrates requirement for directional motility in sand fly infections. PLOS Pathogens 15, e1007828.

Benson, D. A., Karsch-Mizrachi, I., Lipman, D. J., Ostell, J. and Wheeler, D. L. (2008). GenBank. Nucleic Acids Res 36, D25–30.

Billington, K., Halliday, C., Madden, R., Dyer, P., Barker, A. R., Moreira-Leite, F. F., Carrington, M., Vaughan, S., Hertz-Fowler, C., Dean, S., et al. (2023). Genome-wide subcellular protein map for the flagellate parasite Trypanosoma brucei. Nat Microbiol 8, 533–547.

Branche, C., Kohl, L., Toutirais, G., Buisson, J., Cosson, J. and Bastin, P. (2006). Conserved and specific functions of axoneme components in trypanosome motility. J. Cell. Sci. 119, 3443– 3455.

Broadhead, R., Dawe, H. R., Farr, H., Griffiths, S., Hart, S. R., Portman, N., Shaw, M. K., Ginger, M. L., Gaskell, S. J., McKean, P. G., et al. (2006). Flagellar motility is required for the viability of the bloodstream trypanosome. Nature 440, 224–227.

Brun, R. and Schönenberger, null (1979). Cultivation and in vitro cloning or procyclic culture forms of Trypanosoma brucei in a semi-defined medium. Short communication. Acta Trop. 36, 289–292.

de Souza, W. and Souto-Padrón, T. (1980). The paraxial structure of the flagellum of trypanosomatidae. J. Parasitol. 66, 229–236.

Dean, S., Sunter, J., Wheeler, R. J., Hodkinson, I., Gluenz, E. and Gull, K. (2015). A toolkit enabling efficient, scalable and reproducible gene tagging in trypanosomatids. Open Biol 5,.

Deflorin, J., Rudolf, M. and Seebeck, T. (1994). The major components of the paraflagellar rod of Trypanosoma brucei are two similar, but distinct proteins which are encoded by two different gene loci. J Biol Chem 269, 28745–28751.

Eddy, E. M., Toshimori, K. and O’Brien, D. A. (2003). Fibrous sheath of mammalian spermatozoa. Microsc. Res. Tech. 61, 103–115.

Farina, M., Attias, M., Souto-Padron, T. and Souza, W. D. (1986). Further Studies on the Organization of the Paraxial Rod of Trypanosomatids1. The Journal of Protozoology 33, 552– 557.

Ginger, M. L. (2005). Trypanosomatid biology and euglenozoan evolution: new insights and shifting paradigms revealed through genome sequencing. Protist 156, 377–392.

Ginger, M. L., Collingridge, P. W., Brown, R. W. B., Sproat, R., Shaw, M. K. and Gull, K. (2013). Calmodulin is required for paraflagellar rod assembly and flagellum-cell body attachment in trypanosomes. Protist 164, 528–540.

Griffiths, S., Portman, N., Taylor, P. R., Gordon, S., Ginger, M. L. and Gull, K. (2007). RNA interference mutant induction in vivo demonstrates the essential nature of trypanosome flagellar function during mammalian infection. Eukaryotic Cell 6, 1248–1250.

Höög, J. L., Lacomble, S., O’Toole, E. T., Hoenger, A., McIntosh, J. R. and Gull, K. (2014). Modes of flagellar assembly in Chlamydomonas reinhardtii and Trypanosoma brucei. Elife 3, e01479.

Huff, J., Bergter, A., Birkenbeil, J., Kleppe, I., Engelmann, R. and Krzic, U. (2017). The new 2D Superresolution mode for ZEISS Airyscan. Nat Methods 14, 1223–1223.

Hughes, L. C., Ralston, K. S., Hill, K. L. and Zhou, Z. H. (2012). Three-Dimensional Structure of the Trypanosome Flagellum Suggests that the Paraflagellar Rod Functions as a Biomechanical Spring. PLOS ONE 7, e25700.

Hyams, J. S. (1982). The Euglena paraflagellar rod: structure, relationship to other flagellar components and preliminary biochemical characterization. Journal of Cell Science 55, 199– 210.

Inoue, M., Nakamura, Y., Yasuda, K., Yasaka, N., Hara, T., Schnaufer, A., Stuart, K. and Fukuma, T. (2005). The 14-3-3 proteins of Trypanosoma brucei function in motility, cytokinesis, and cell cycle. J Biol Chem 280, 14085–14096.

Irons, M. J. and Clermont, Y. (1982a). Formation of the outer dense fibers during spermiogenesis in the rat. Anat. Rec. 202, 463–471.

Irons, M. J. and Clermont, Y. (1982b). Kinetics of fibrous sheath formation in the rat spermatid. Am. J. Anat. 165, 121–130.

Kohl, L., Sherwin, T. and Gull, K. (1999). Assembly of the paraflagellar rod and the flagellum attachment zone complex during the Trypanosoma brucei cell cycle. J. Eukaryot. Microbiol. 46, 105–109.

Lacomble, S., Portman, N. and Gull, K. (2009). A protein-protein interaction map of the Trypanosoma brucei paraflagellar rod. PLoS ONE 4, e7685.

Maga, J. A., Sherwin, T., Francis, S., Gull, K. and LeBowitz, J. H. (1999). Genetic dissection of the Leishmania paraflagellar rod, a unique flagellar cytoskeleton structure. J. Cell. Sci. 112 (Pt 16), 2753–2763.

Miki, K., Willis, W. D., Brown, P. R., Goulding, E. H., Fulcher, K. D. and Eddy, E. M. (2002). Targeted disruption of the Akap4 gene causes defects in sperm flagellum and motility. Dev. Biol. 248, 331–342.

Moran, J., McKean, P. G. and Ginger, M. L. (2014). Eukaryotic Flagella: Variations in Form, Function, and Composition during Evolution. BioScience 64, 1103–1114.

Moreira-Leite, F. F., de Souza, W. and da Cunha-e-Silva, N. L. (1999). Purification of the paraflagellar rod of the trypanosomatid Herpetomonas megaseliae and identification of some of its minor components. Mol Biochem Parasitol 104, 131–140.

Petersen, C., Füzesi, L. and Hoyer-Fender, S. (1999). Outer dense fibre proteins from human sperm tail: molecular cloning and expression analyses of two cDNA transcripts encoding proteins of approximately 70 kDa. Mol. Hum. Reprod. 5, 627–635.

Portman, N. and Gull, K. (2010). The paraflagellar rod of kinetoplastid parasites: from structure to components and function. Int. J. Parasitol. 40, 135–148.

Portman, N., Lacomble, S., Thomas, B., McKean, P. G. and Gull, K. (2009). Combining RNA interference mutants and comparative proteomics to identify protein components and dependences in a eukaryotic flagellum. J. Biol. Chem. 284, 5610–5619.

Pullen, T. J., Ginger, M. L., Gaskell, S. J. and Gull, K. (2004). Protein targeting of an unusual, evolutionarily conserved adenylate kinase to a eukaryotic flagellum. Mol. Biol. Cell 15, 3257– 3265.

Rascón, A., Soderling, S. H., Schaefer, J. B. and Beavo, J. A. (2002). Cloning and characterization of a cAMP-specific phosphodiesterase (TbPDE2B) from Trypanosoma brucei. Proc Natl Acad Sci U S A 99, 4714–4719.

Redmond, S., Vadivelu, J. and Field, M. C. (2003). RNAit: an automated web-based tool for the selection of RNAi targets in Trypanosoma brucei. Mol Biochem Parasitol 128, 115–118.

Ridgley, E., Webster, P., Patton, C. and Ruben, L. (2000). Calmodulin-binding properties of the paraflagellar rod complex from Trypanosoma brucei. Mol Biochem Parasitol 109, 195–201.

Santrich, C., Moore, L., Sherwin, T., Bastin, P., Brokaw, C., Gull, K. and LeBowitz, J. H. (1997). A motility function for the paraflagellar rod of Leishmania parasites revealed by PFR-2 gene knockouts. Mol. Biochem. Parasitol. 90, 95–109.

Schindelin, J., Arganda-Carreras, I., Frise, E., Kaynig, V., Longair, M., Pietzsch, T., Preibisch, S., Rueden, C., Saalfeld, S., Schmid, B., et al. (2012). Fiji: an open-source platform for biological-image analysis. Nat Methods 9, 676–682.

Schmelling, N. M., Lehmann, R., Chaudhury, P., Beck, C., Albers, S.-V., Axmann, I. M. and Wiegard, A. (2017). Minimal tool set for a prokaryotic circadian clock. BMC Evol Biol 17, 169.

Shaw, S., DeMarco, S. F., Rehmann, R., Wenzler, T., Florini, F., Roditi, I. and Hill, K. L. (2019). Flagellar cAMP signaling controls trypanosome progression through host tissues. Nat Commun 10, 803.

Subota, I., Julkowska, D., Vincensini, L., Reeg, N., Buisson, J., Blisnick, T., Huet, D., Perrot, S., Santi-Rocca, J., Duchateau, M., et al. (2014). Proteomic analysis of intact flagella of procyclic Trypanosoma brucei cells identifies novel flagellar proteins with unique sub-localization and dynamics. Mol. Cell Proteomics 13, 1769–1786.

Sunter, J. D. and Gull, K. (2016). The Flagellum Attachment Zone: “The Cellular Ruler” of Trypanosome Morphology. Trends Parasitol. 32, 309–324.

Tarnasky, H., Cheng, M., Ou, Y., Thundathil, J. C., Oko, R. and van der Hoorn, F. A. (2010). Gene trap mutation of murine outer dense fiber protein-2 gene can result in sperm tail abnormalities in mice with high percentage chimaerism. BMC Dev. Biol. 10, 67.

Wheeler, R. J. (2017). Use of chiral cell shape to ensure highly directional swimming in trypanosomes. PLoS Comput. Biol. 13, e1005353.

Woods, A., Sherwin, T., Sasse, R., MacRae, T. H., Baines, A. J. and Gull, K. (1989). Definition of individual components within the cytoskeleton of Trypanosoma brucei by a library of monoclonal antibodies. J. Cell. Sci. 93 (Pt 3), 491–500.

Woodward, R., Carden, M. J. and Gull, K. (1994). Molecular characterisation of a novel, repetitive protein of the paraflagellar rod in Trypanosoma brucei. Mol. Biochem. Parasitol. 67, 31–39.

Woolley, D. M. (1971). Striations in the peripheral fibers of rat and mouse spermatozoa. J. Cell Biol. 49, 936–939.

Yubuki, N., Huang, S. S. C. and Leander, B. S. (2016). Comparative Ultrastructure of Fornicate Excavates, Including a Novel Free-living Relative of Diplomonads: Aduncisulcus paluster gen. et sp. nov. Protist 167, 584–596.

Zhang, J., Wang, H., Imhof, S., Zhou, X., Liao, S., Atanasov, I., Hui, W. H., Hill, K. L. and Zhou, Z. H. (2021). Structure of the trypanosome paraflagellar rod and insights into non-planar motility of eukaryotic cells. Cell Discov 7, 51.

Zhao, W., Li, Z., Ping, P., Wang, G., Yuan, X. and Sun, F. (2018). Outer dense fibers stabilize the axoneme to maintain sperm motility. J. Cell. Mol. Med. 22, 1755–1768.

