## Supplementary figures for "Dramatic reduction in trypanosome motility occurs without large-scale changes to paraflagellar rod ultrastructure"

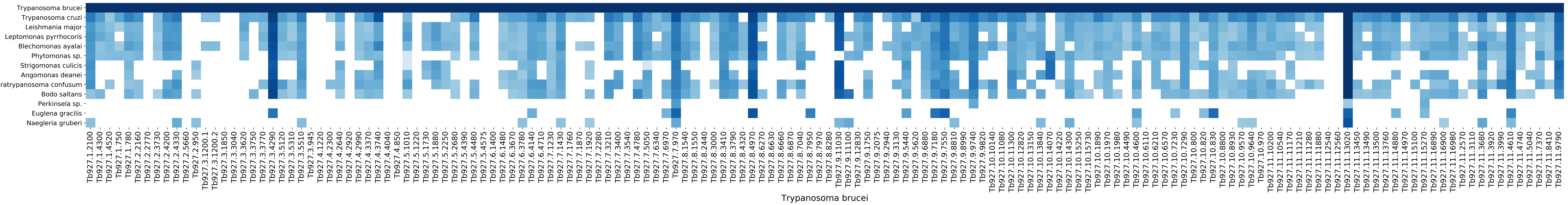

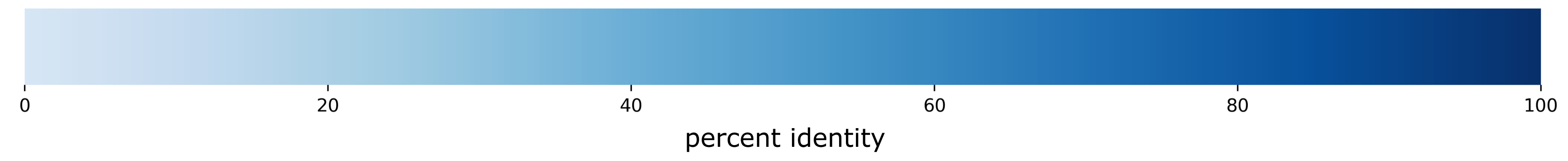


**Figure S1.** Heatmap of the reciprocal best BLAST analysis results. The colour indicates the percent identity between the *T. brucei* PFR protein and the ortholog in that species. White means no ortholog present.

A

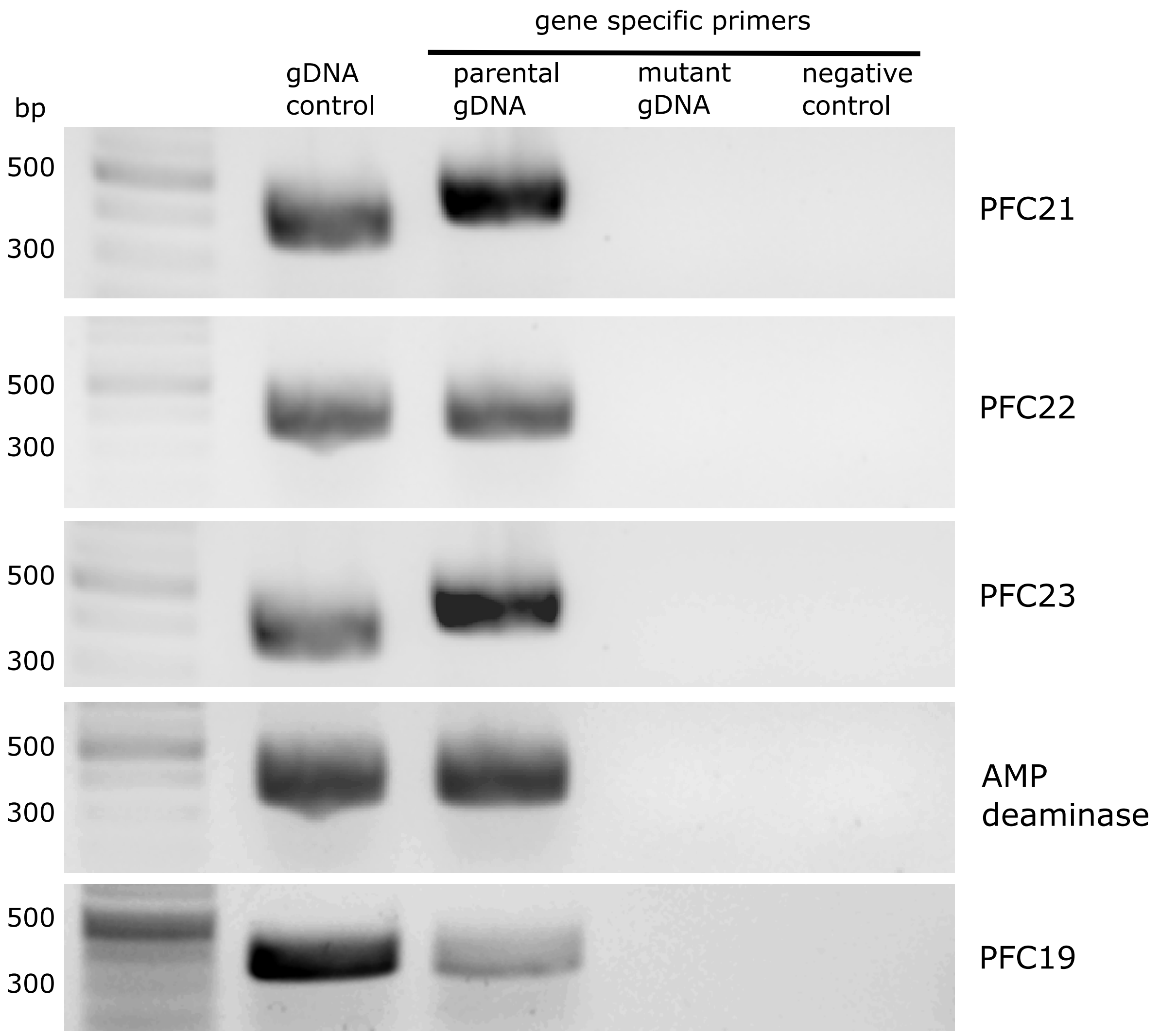


B

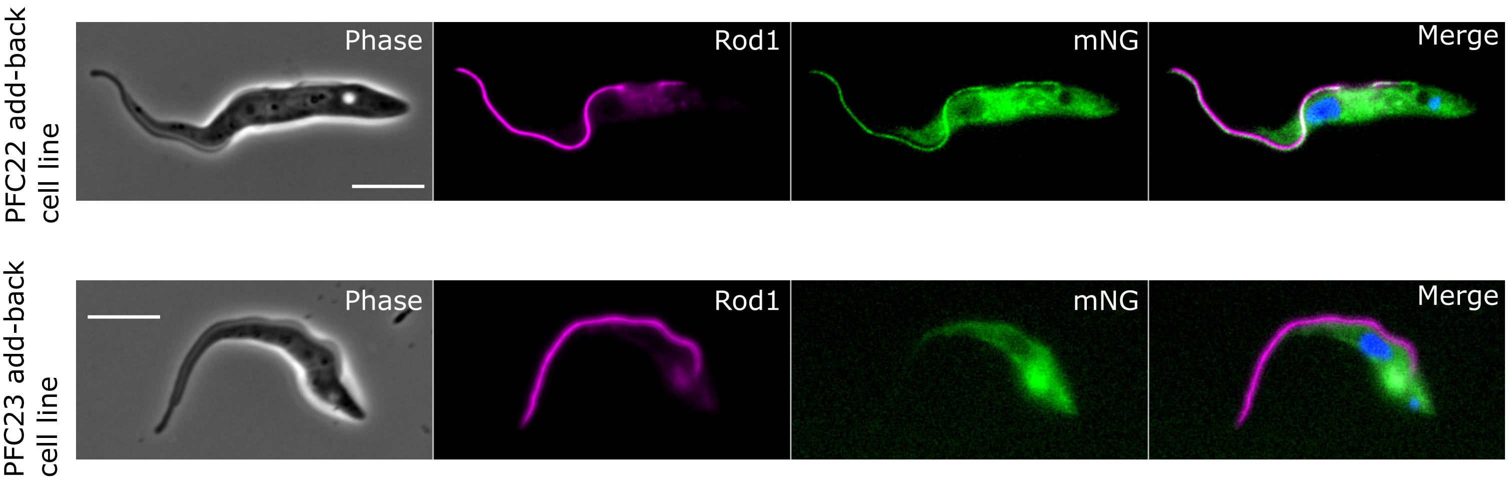


Figure S2. A) PCR analysis of knockout mutants. Loss of deleted gene analysed using gene specific primers. Gene specific primers were tested against parental genomic DNA. Negative control was water only. B) Example images of PFC22 and PFC23 add back cells expressing Rod1 endogenously tagged with mCherry and PFC22 and PFC23 tagged with mNG respectively. Far left is a phase image, followed by the Rod1 marker and then PFC22 or PFC23. Merge shows mCherry, mNG and DNA stained with Hoechst 33342 (blue). Scale is 5 μm.

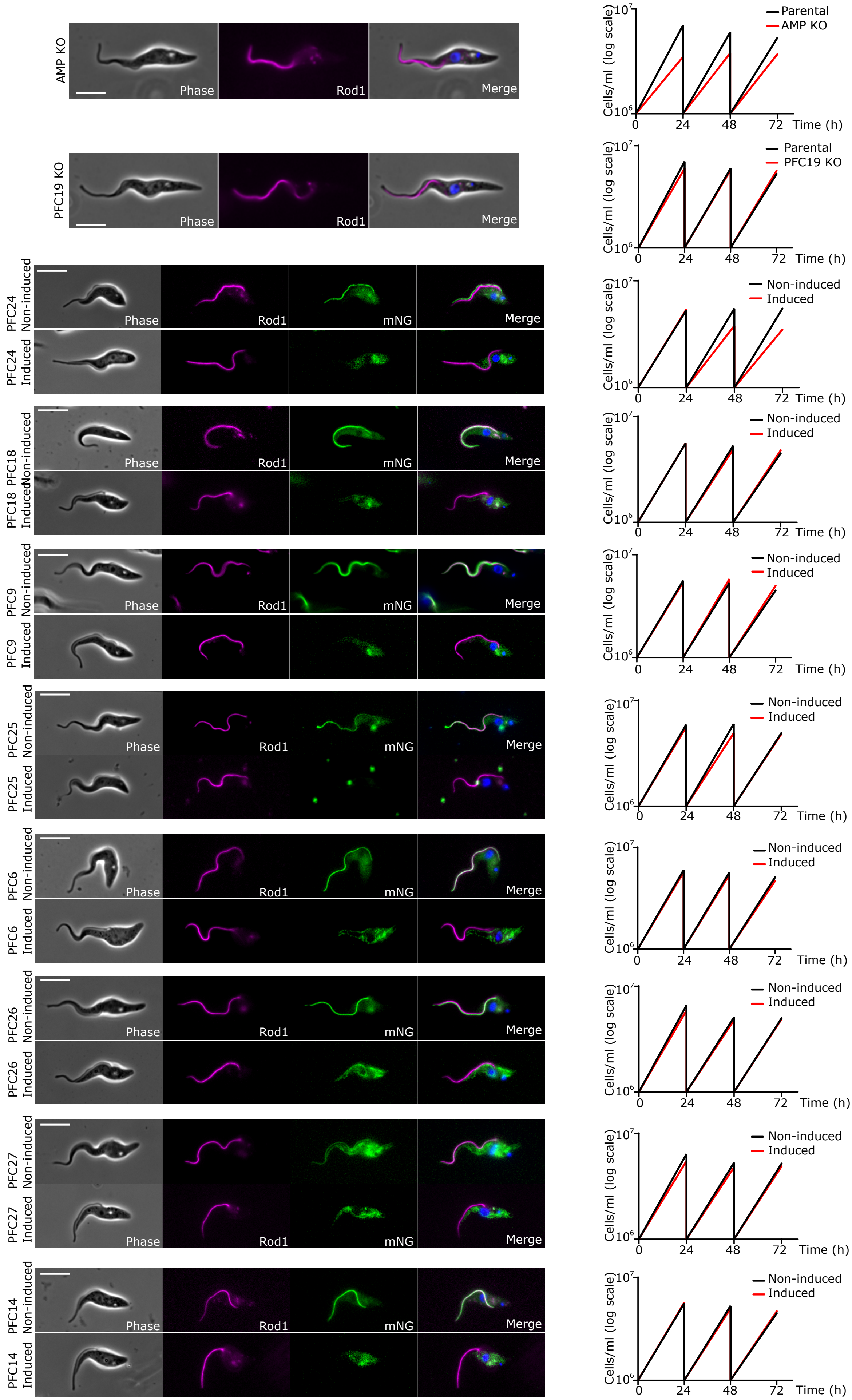


Figure S3. Example images of PFC19 and AMP deaminase deletion mutants expressing Rod1 tagged with mCherry. Left is a phase image, followed by the Rod1 marker. Merge shows phase, mCherryand DNA stained with Hoechst 33342 (blue). Scale is 5 µm. Growth of parental and KO cells was followed for 72 h. Example images of cells before and after RNAi induction for 72 hours with doxycycline (1 µg/ml). Cells express Rod1 endogenously tagged with mCherry and PFR protein of interest tagged, with mNG to monitor RNAi mediated protein knockdown. Far left is a phase image, followed by the Rod1 marker and then the PFR protein. Merge shows phase, mCherry, mNG and DNA stained with Hoechst 33342 (blue). Growth of non-induced and induced cells was followed for 72 hours.

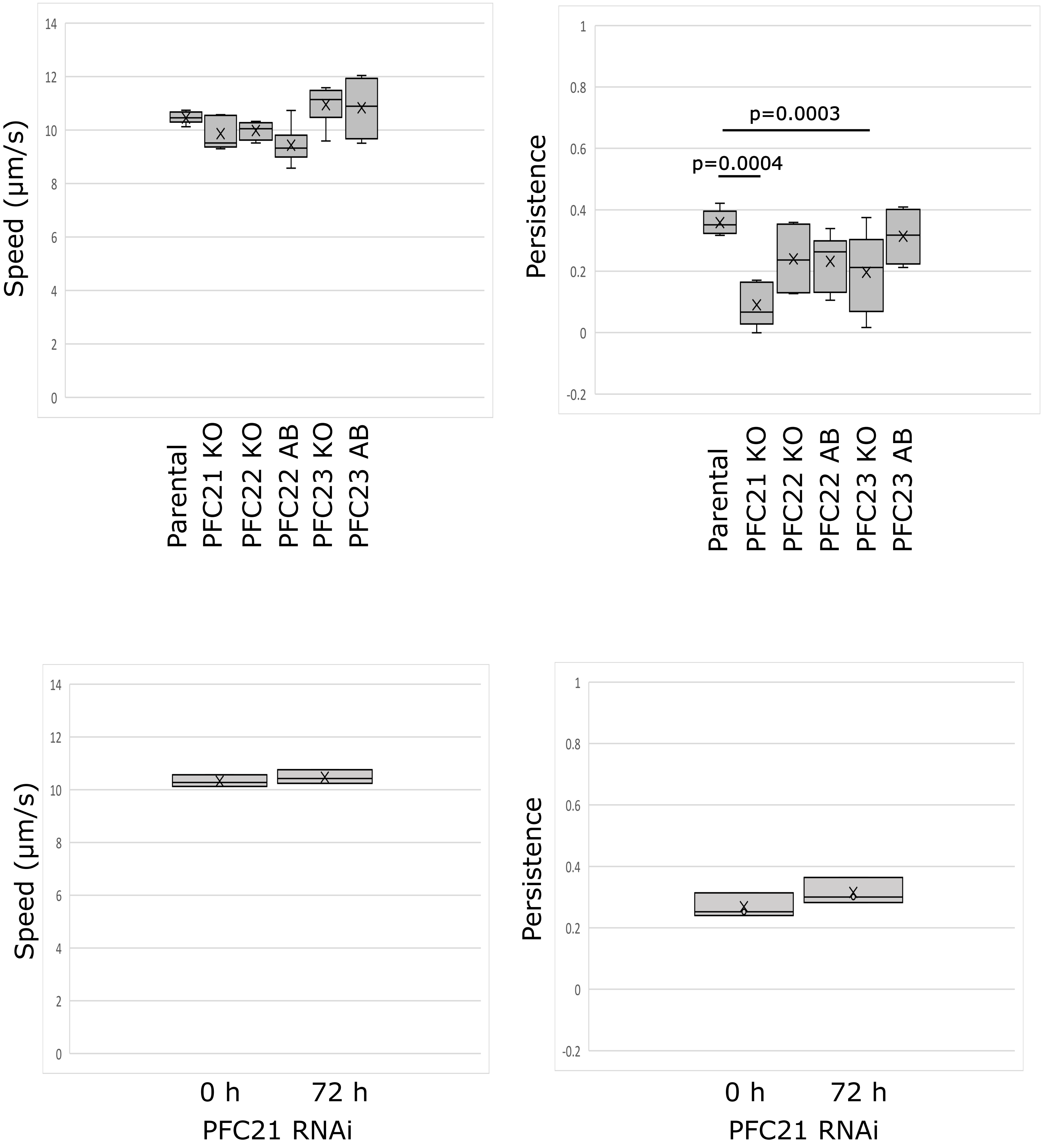


Figure S4. Motility analysis of PFC mutant cell lines. Box and whisker plots of mean speed and mean directional persistence showing the minimum value, first quartile, median, third quartile and maximum value. The black X is the mean value. Each individual motility assay measures the speed and persistence of >700 cells. The mean from each of these assays for speed and directional persistence is then calculated and plotted here. For each cell line the motility assay was performed 3-6 times. P-value was calculated using a 2-tailed t-test.
